# CST–Polymeraseα-primase solves a second telomere end-replication problem

**DOI:** 10.1101/2023.10.26.564248

**Authors:** Hiroyuki Takai, Valentina Aria, Pamela Borges, Joseph T. P. Yeeles, Titia de Lange

**Affiliations:** Laboratory for Cell Biology and Genetics, Rockefeller University, New York, USA; Medical Research Council Laboratory of Molecular Biology, Cambridge, CB2, 0QH

## Abstract

Telomerase adds G-rich telomeric repeats to the 3ʹ ends of telomeres^1^, counteracting telomere shortening caused by loss of telomeric 3ʹ overhangs during leading-strand DNA synthesis (“the end-replication problem”^2^). We report a second end-replication problem that originates from the incomplete duplication of the C-rich telomeric repeat strand by lagging-strand synthesis. This problem is solved by CST–Polymeraseα(Polα)-primase fill-in synthesis. *In vitro,* priming for lagging-strand DNA replication does not occur on the 3’ overhang and lagging-strand synthesis stops in an ∼150-nt zone more than 26 nt from the end of the template. Consistent with the *in vitro* data, lagging-end telomeres of cells lacking CST–Polα-primase lost ∼50-60 nt of CCCTAA repeats per population doubling (PD). The C-strands of leading-end telomeres shortened by ∼100 nt/PD, reflecting the generation of 3’ overhangs through resection. The measured overall C-strand shortening in absence of CST–Polα-primase fill-in is consistent with the combined effects of incomplete lagging-strand synthesis and 5ʹ resection at the leading-ends. We conclude that canonical DNA replication creates two telomere end-replication problems that require telomerase to maintain the G-strand and CST–Polα-primase to maintain the C-strand.

## Main

Human telomeres are composed of long arrays of duplex TTAGGG repeats ending in a 3ʹ overhang of the G-rich strand (Fig. 1a). The 3ʹ overhangs allow telomeres to adopt the protective t-loop configuration^3,4^ and serve as primers for telomerase. During the replication of telomeres, the C-rich 5ʹ-ended strand is the template for leading-strand DNA synthesis while the TTAGGG repeat strand templates lagging-strand DNA synthesis (Fig. 1a). The end-replication problem was originally proposed to involve removal of the RNA primer of the most terminal Okazaki fragment, predicting cumulative sequence loss at the 5ʹ ends of lagging-strand products^5,6^ (Extended Data Figure 1a). Telomerase was initially proposed to solve this problem by elongating the G-rich strand before DNA replication^7^, thereby extending the template for Okazaki fragment synthesis (Extended Data Figure 1b). However, once it had become clear that telomeres of most organisms carry constitutive 3ʹ overhangs, which cannot be regenerated by leading-strand DNA synthesis, a new version of the end-replication problem emerged^2^ (Fig. 1a; Extended Data Figure 1c). In this version, the sequence loss –equivalent to the length of the 3ʹ overhang– occurs on the leading-strand and is counteracted by telomerase adding a 3ʹ overhang to the leading-strand product^4^ (Fig. 1a). In agreement, human telomerase was shown to act after DNA replication, extending the 3ʹ ends of both leading- and lagging-strand DNA synthesis products^8,9^. Ini)al 5ʹ end resection of the leading-strand products, presumed to be blunt, is needed for their extension by telomerase and serves to regenerate the 3ʹ overhangs in cells lacking telomerase^10–13^ (Fig. 1a). According to this scenario, telomerase solves the leading-strand end-replication problem^2,13^ while the constitutive 3ʹ overhangs solve the lagging-strand problem (Fig. 1a). In agreement with the idea that lagging-strand synthesis can initiate on the 3ʹ overhang, RNA primers were observed along the 3ʹ overhangs of human telomeres^11^ (Fig. 1a; Extended Data Fig. 1). However, whether these priming events resulted from canonical lagging-strand DNA replication was not established. The data described below argue that the measured priming events reflected CST–Polα-primase fill-in synthesis. Here we use an *in vitro* DNA replication system to determine priming sites for the last Okazaki fragments generated by the replisome and identify an additional end-replication problem that it is solved by CST–Polα-primase *in vivo*.

**Figure 1.**
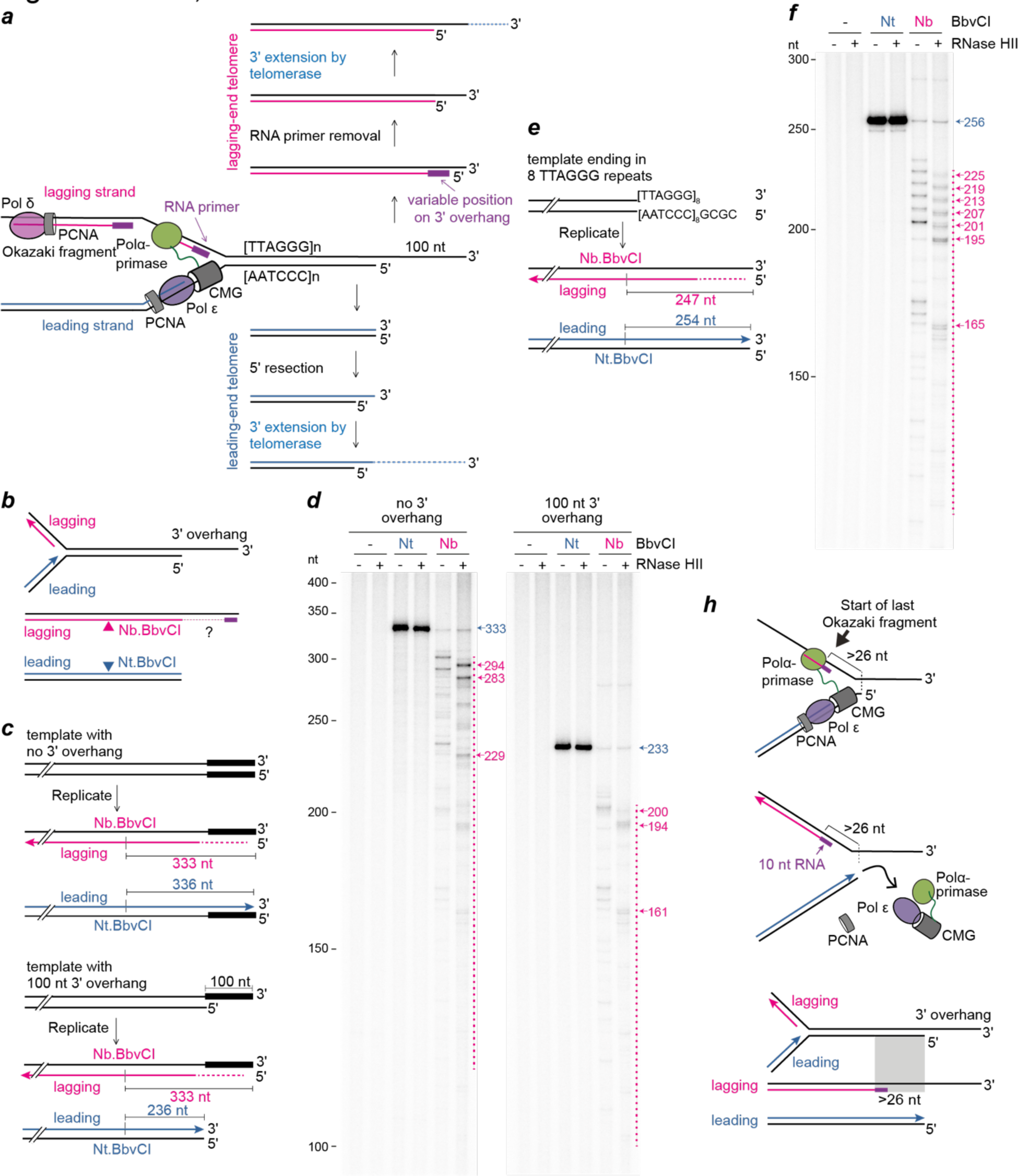
In vitro replication of linear templates reveals a second end-replication problem. **a**, Schematic of the end-replication problem at a telomere with a 100 nt 3ʹ overhang and its solution by telomerase prior to this study (modified from **refs. 7 and 10**^7,10^). **b**, Schematic of the in vitro replication approach to determine the products formed by the replisome at DNA ends. **c**, Diagrams of replication templates and anticipated leading and lagging-strand products. The location of the BbvCI site and the anticipated products of nicking by Nb.BbvCI and Nt.BbvCI are shown. **d**, Denaturing 5.5% polyacrylamide/urea gel analysis of a 30 min replication reaction performed on the templates illustrated in (**c**). Products were post-replicatively digested with Nt./Nb.BbvCI and RNase HII as indicated. The length in nucleotides of major reaction products, derived from the plots in Extended Data Fig. 2b, are shown. The distribution of Nb.BbvCI-dependent products is marked by a dashed line. **e**, Diagram of a replication template containing 8 terminal TTAGGG repeats and anticipated leading and lagging-strand products after digestion with Nb.BbvCI and Nt.BbvCI. **f**, Denaturing 5.2% polyacrylamide/urea gel analysis of a replication reaction performed on the template illustrated in **e**, analyzed as in **d**. **h**, Schematic illustrating that the last Okazaki fragment (after primer removal) starts >26 nt from the 5ʹ end of the template and the terminal structures predicted to be created by replication of a linear DNA. To enable clear visualization of the Nb.BbvCI products, the Nt.BbvCI product bands are saturated in (**d**) and (**f**).

To determine how the replisome behaves at the end of a linear DNA template, we utilized an *in vitro* reconstituted DNA replication system to map the ends of nascent lagging- and leading-strands. In this system, replisomes are assembled with purified *Saccharomyces cerevisiae* proteins that perform complete leading- and lagging-strand replication at the *in vivo* rate^15,16^. Origin-dependent DNA replication was performed on linear templates that either carried a 100 nt 3ʹ overhang or the same DNA sequence in duplex form (no overhang) (Fig. 1b,c and Extended Data Fig. 2a). Prior to analysis through denaturing polyacrylamide gels, the radiolabelled replication products were nicked with Nb.BbvCI or Nt.BbvCI which liberate nascent lagging- and leading-strand ends, respectively (Fig. 1b,c). Reaction products digested with Nt.BbvCI were insensitive to RNase HII and migrated as single bands of close to the expected lengths for fully-extended replication products (236 and 336 nt for templates with and without overhang, respectively) (Fig.1d), establishing that leading-strand DNA synthesis efficiently completes the replication of linear DNA templates, as predicted by data on yeast telomere replication^17,18^. In contrast, diges)on of replica)on products with Nb.BbvCI generated ladders of bands that varied in intensity and were sensi)ve to RNase HII treatment, confirming that they corresponded to the 5ʹ ends of nascent lagging strands. The shortest RNase HII sensitive products were ∼150-200 nucleotides shorter than the maximum theoretical Nb.BbvCI cleavage product (Fig.1d and Extended Data Fig. 2b). Strikingly, after RNase HII digestion, the longest RNase HII sensitive products were ∼30 nt shorter than the corresponding leading-strand products regardless of the presence of a 3ʹ overhang (Fig. 1d and Extended Data Fig. 2b). These results reveal that the core eukaryotic replisome initiates the final Okazaki fragment at multiple positions within ∼150-200 nucleotides of the end of a linear template. Additionally, they show that the replisome is unable to initiate new Okazaki fragments once it gets to within ∼20 nt of the end of the leading-strand template and consequently it cannot sustain canonical lagging-strand synthesis along a 3ʹ overhang.

To mimic the replication of telomere ends, we generated a linear template with eight TTAGGG repeats at the 3ʹ end of the lagging-strand template (Fig. 1e). Replication of this template produced 6 evenly spaced RNase HII sensitive products (Fig.1f and Extended Data Fig. 2c). The lengths of these products indicate that they represent Okazaki fragments initiated within the telomeric repeats using ATP to begin primer synthesis (Extended Data Fig. 2d). Notably, the presence of 6, rather than 8, bands suggests that the replisome is unable to utilize the final 16 nt of TTAGGG repeat DNA to initiate new Okazaki fragments (Extended Data Fig. 2d), even when the concentration of Polα-primase in the reaction was increased 4-fold (Extended Data Fig. 2e,f). To further validate this conclusion, we generated a template with an identical sequence but with 121 bp of dsDNA downstream of the 8 TTAGGG repeats and observed 8 evenly spaced RNase HII sensitive products, demonstrating that Okazaki fragment synthesis was initiated on each of the eight repeats (Extended Data Fig. 3a-c). Collectively, these data demonstrate that for terminal TTAGGG repeats, it is the proximity to the end of the template that renders them incompatible with Okazaki fragment initiation. This behaviour is consistent with recent cryo-EM structures of yeast and human replisomes containing Polα-primase^19^. In both structures, the primase active site is positioned ∼75 Å from the point of template unwinding (Extended Data Fig. 3d), which equates to ∼20 nucleotides of ssDNA. Consequently, in this configuration, it is unlikely that Polα-primase can prime within the last ∼20 nt of a linear template because the parental DNA duplex will be fully unwound before the template has reached the primase active site. Moreover, because the positioning of Polα-primase in the replisome is highly conserved between budding yeast and human (Extended Data Fig. 3d), it is reasonable to expect that lagging-strand replication by human replisomes will behave the same as budding yeast replisomes during the duplication of telomeres. Due to this additional end-replication problem, the C-rich telomeric strand is predicted to continuously shorten by >26 nt and this shortening will not be mitigated by extension of the G-rich strand by telomerase.

Sequence loss from the 5ʹ end of the C-rich telomeric strand can be counteracted by Polα-primase in association with the CST (Ctc1, Stn1, Ten1) complex^20,21^. CST is an RPA-like ssDNA binding protein that interacts with Polα-primase and stimulates its activity^22–31^. At mammalian telomeres, CST–Polα-primase can fill in telomere ends that have been hyper-resected, thereby preventing excessive telomere shortening^10,11,32,33^. Prior work has shown that the C-rich strand progressively shortens when CST–Polα-primase fill-in is disabled^32,33^ but the rate of shortening at leading- and lagging-end telomeres and the cause of shortening remained unknown.

We set out to determine whether lagging-strand DNA synthesis results in shortened C-strands *in vivo*. CsCl-gradients can be used to separate telomeres generated by leading- and lagging-strand DNA synthesis based on their differential incorporation of BrdU^14^ (Extended Data Fig. 4a). This method yields limited amounts of DNA, necessitating quantification of the 3ʹ overhangs by the sensitive in-gel hybridization method^10,34^. We applied this approach to CTC1^F/F^ HCT116 colon carcinoma cells^32^ from which *CTC1* can be deleted upon induction of Cre with tamoxifen (4-OHT). CTC1^F/F^ cells express telomerase but were rendered telomerase-negative through bulk CRISPR/Cas9 KO of hTERT and further inhibition of telomerase activity with BIBR1532^35^ so that no extension of the 3ʹ overhangs by telomerase will take place (Fig. 2a; Extended Data Fig. 4b,c). The relative 3ʹ overhang signals from six independent experiments showed that deletion of *CTC1* induced a substantial increase in the relative 3ʹ overhang signals at both leading- and lagging-end telomeres, as expected from prior work^10,11,32,33^ (Fig. 2b,c).

**Figure 2.**
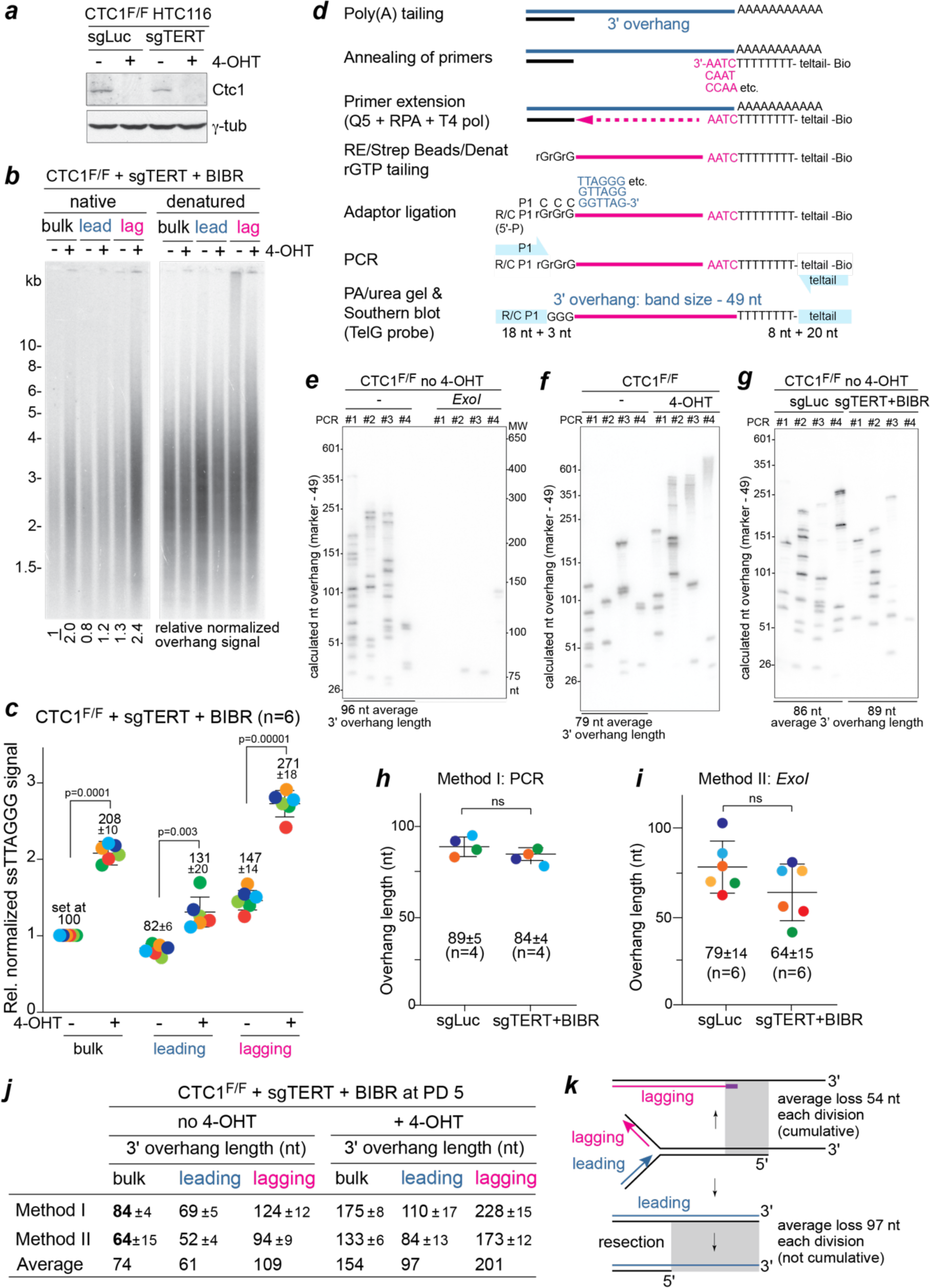
C-strand loss at leading- and lagging-end telomeres in absence of CST. ***a***, Immunoblot for Ctc1 in CTC1^F/F^ HCT116 cells (plus and minus 4-OHT to induce Cre) after bulk CRISPR/Cas9 knockout of hTERT (sgTERT). sgLuc: Luciferase control. ***b***, In-gel overhang assay for relative overhang signals of leading-(lead) and lagging-(lag) end telomeres of telomerase-deficient CTC1^F/F^ cells isolated on CsCl gradients. Left: hybridization with TelC to detect the G-strand overhang. Right: same gel reprobed with TelC after in situ denaturation, detecting total telomeric DNA for normalization (normalized values relative to lane 1). Cells were harvested at PD 5 after CTC1 deletion. ***c***, Graph showing the relative overhang signals and SDs from six independent experiments as in (***b***). P values determined by unpaired t-test. ***d***, Schematic of the PCR assay for telomeric overhang length. See text and methods for details. ***e***, PCR overhang assays (four PCRs per DNA sample) on genomic DNA with and without pre-treatment with *E. coli Exo*I showing the specificity of the assay for 3ʹ overhangs. Calculated 3ʹ overhang lengths are on the left. Average sizes of 3ʹ overhangs are given below the panel. ***f***, PCR overhang assay detects elongated 3ʹ telomeric overhang in cells lacking Ctc1 as in (***e***). The accuracy of the assay for long overhangs has not been determined. ***g***, PCR overhang assay on DNA from CTC1^F/F^ cells (no 4-OHT) with and without telomerase activity as in (***e***). ***h***, Graph showing average 3ʹ overhang lengths and SDs from 4 biological replicates (4 PCRs per replicate) as in (***g***). ns, not significant based on unpaired t-test. ***i***, Graph showing average 3ʹ overhang lengths and SDs determined by Method II (Extended Data Fig. 6) on 6 biological replicates. ***j***, Absolute overhang lengths at leading- and lagging-end telomeres in telomerase-deficient cells with and without Ctc1. The values in bold from panels (***h***) and (***i***) were used to calibrate the relative overhang values in (***c***)*. **k**,* C-strand loss in telomerase-negative cells lacking Ctc1 based on (***j***) and calculations given in Extended Data Fig. 7.

Because the in-gel detection of telomeric overhangs yields only relative values, we aimed to determine the absolute length of the 3ʹ overhang in at least one of the samples to calibrate the relative overhang values from the in-gel assays. We developed two methods to measure absolute lengths of the 3ʹ overhang in cells containing CST in the presence or absence of telomerase activity. The first method, inspired by primer extension experiments performed with yeast and human telomeric DNA^34,36^, is based on a PCR amplification of a fill-in product generated with poly(A) tailed 3ʹ overhangs as a template (Fig. 2d). This method relies on annealing the poly(A)-tailed telomeres to a set of oligodT primers that represent the six possible 3ʹ end sequences of telomeres followed by primer extension by bacterial DNA polymerases that lack strand-displacement activity. The oligodT primers also contained a specific sequence for PCR (teltail) and a 5ʹ Biotin for isolation of the extension products. The extension products are tailed with rGTP and ligated to an adapter allowing generation of PCR products that can be analysed on blots of polyacrylamide/urea gels hybridized to a radiolabelled G-strand oligonucleotide. The method was tested on a telomere model substrate featuring a stretch of duplex telomeric repeats and a [TTAGGG]n 3ʹ overhang of 56 nt (Extended Data Fig. 5a,b). This substrate yielded a PCR product of the expected length that was not observed when the 3ʹ overhang was first removed with the *E. coli* 3ʹ exonuclease *Exo*I (Extended Data Fig. 5b).

DNA from Ctc1-proficient cells yielded sets of PCR products that were largely absent when the DNA was first treated with *Exo*I to remove the 3ʹ overhangs (Fig. 2e). As is the case for the PCR products in the Single Telomere Length Analysis (STELA^37^) assay, parallel PCRs on the same DNA sample yielded different sets of bands, indicating that each band likely represents the 3ʹ overhang of a single telomere. As expected, the PCR products from cells lacking Ctc1 were substantially longer (Fig. 2f). Control experiments with oligos designed to represent fill-in products from 3ʹ overhangs with 5-15 repeats indicated that the assay was robust, yielding products with the expected length and intensity (Extended Data Fig. 5c,d). An oligo containing 20 repeats was also readily detected although with greater variation in intensity (Extended Data Fig. 5d), indicating that some of the products obtained from longer 3’ overhangs could escape detection. Using this approach, the average lengths of the 3ʹ overhangs in genomic DNAs from CTC1^F/F^ and CTC1^F/F^ sgTERT cells treated with BIBR1532 (but not with 4-OHT) was determined in four independent experiments (Fig. 2g). The average overhang length of each DNA sample was derived by summing the sizes of all bands observed in four PCRs, dividing the sum by the number of detected bands, and subtracting the 49 nt non-telomeric sequences in the products (Fig. 2d-g). The average 3ʹ overhang lengths in cells with and without telomerase were 89 and 84 nt, respectively (Fig. 2h). The similarly sized 3ʹ overhangs in cells with and without telomerase is consistent with data from in-gel hybridization experiments which showed a minimal difference in the relative 3ʹ overhang signals (Extended Data Fig. 5e).

In an orthogonal approach to determine the absolute 3ʹ overhang lengths, we measured the shortening of the G-rich telomeric strand upon digestion of duplex DNA with *Exo*I. This method is reminiscent of experiments done on yeast telomeres, which are sufficiently short to be analyzed by PCR before and after *Exo*I treatment^17^. To analyze the longer telomeres of HCT116 cells, genomic DNA was treated with *Exo*I, digested with *Mbo*I and *Alu*I to generate telomeric restriction fragments, and the reduction in the size of the telomeric G-strand was measured on alkaline agarose gels by in-gel hybridization (Extended Data Fig. 6). This method was tested on the model telomere with a 3ʹ overhang of defined length (Extended Data Fig. 6a). *Exo*I digestion of the linear model telomere bearing a 3ʹ overhang reduced the size of the G-strand by 57 nt which is close to the expected value of 56 nt (Extended Data Fig. 6a). The *Exo*I method was applied to DNA from Ctc1-proficient cells with and without telomerase (Extended Data Fig. 6b-e) which showed 3ʹ overhang lengths of 79 and 64 nt for cells with and without telomerase, respectively (Fig. 2h). These values are close to those obtained by the PCR method.

The overhang lengths determined by the two methods were used to calibrate the relative overhang signals measured on separated leading- and lagging-end telomeres (Fig. 2j). According to this analysis, telomerase-negative cells lacking Ctc1 have 3ʹ overhangs of ∼97 and ∼201 nt at their leading- and lagging-end telomeres, respectively. Taking these values into account, our modelling (Extended Data Fig. 7) indicates that the leading-end telomeres lose 97 nt of C-rich sequences due to resection while the lagging-end telomeres lose ∼54 nt. The finding of this C-strand loss at lagging-end telomeres is consistent with the *in vitro* analysis of the lagging-end replication products. We cannot exclude that the loss of C-strand sequences from the lagging-end telomeres is due to exonucleolytic attack, rather than a lagging-strand replication deficiency. However, we consider this possibility unlikely because it would require a regulatory mechanism that enforces distinct outcomes of resection at the leading- and lagging-end telomeres. Furthermore, a previous study showed that no resection occurred at lagging-end telomeres in human cells^14^, although lagging-end resection was observed in mouse cells^10^. This difference in resection at human and mouse telomeres may be explained based on the recently identified an ATC-5ʹ-P binding site in POT1 proteins^38^ (referred to as the POT-hole). End-binding by the POT-hole could block 5ʹ resection of the last Okazaki fragment, which is predicted to end in ATC-5ʹ-P^18^. While human telomeres have only one POT1 protein that has the POT-hole, mouse telomeres have two POT1 proteins^36^ one of which lacks a POT-hole^35^, possibly leading to less repression of resection lagging-strand mouse telomeres.

Based on the inferred average C-strand shortening caused by resection at the leading-ends (97 nt) and incomplete synthesis at the lagging-ends (54 nt) (Fig. 2j), we predicted how the length of the 3ʹ overhang will change with cell divisions when Ctc1 is absent from telomerase-negative cells (Fig. 3a; Extended Data Fig. 7). We then measured the relative normalized 3ʹ overhang signals in telomerase-negative cells from which *CTC1* was deleted using the in-gel overhang assay (Fig. 3b). The results from three independent experiments showed that the observed relative overhang change was nearly identical to the predicted change (Fig. 3c).

**Figure 3.**
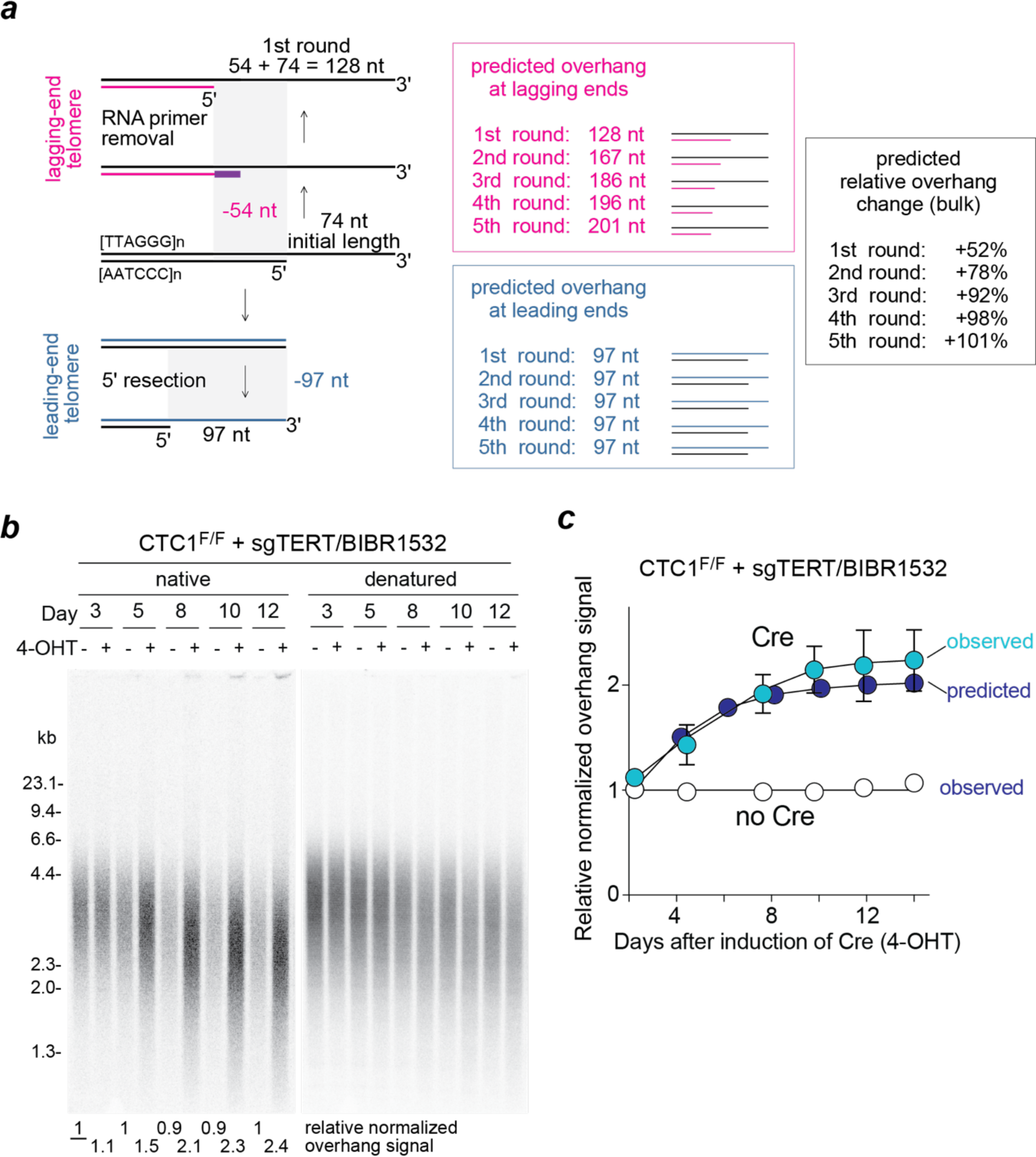
Telomeric overhangs increase at the predicted rate after *CTC1* deletion. *a*, Predicted C-strand loss at leading- and lagging-end telomeres in telomerase- and Ctc1-deficient cells undergoing five cell divisions. The predicted relative overhang change (black) was calculated based on the initial overhang length (74 nt; Fig. 2j) and changes of predicted overhang at lagging- (red) and leading- (green) ends (Fig. 2j, k and Extended Data Fig. 6). ***b***, In-gel overhang assays on telomerase-deficient (sgTERT + BIBR1532) CTC1^F/F^ cells (with or without 4-OHT). Relative normalized overhang signals are given below the gel with the first lane set to 1. ***c***, Graph showing close agreement with the predicted (as in panel **a** black) and observed 3ʹ overhang changes. Values are from 3 independent experiments as in (***b***) with SDs shown.

To further test whether the shortening of telomeric DNA *in vivo* is consistent with the predictions in Fig. 2i, the length changes of G- and C-rich strands were examined after *CTC1* deletion from telomerase-positive cells (Fig. 4a,b; Extended Data Fig. 8a,b). The G-strand was rapidly extended by ∼240 nt/PD when *CTC1* was deleted, consistent with the lack of inhibition of telomerase by CST (Fig. 4a-c; Extended Data Fig. 8a-c; Extended Data Fig. 9). As previously observed^32,33^, the C-strand shortened, while the G-strand elongated, confirming that the replisome is incapable of initiating Okazaki fragment synthesis on the 3ʹ overhang. Based on the inferred rate of C-strand sequence loss at the lagging- and leading-end telomeres, the overall rate of C-strand shortening is predicted to be ∼76 nt/PD, which is close to the observed rate of C-strand shortening (Fig. 4d; Extended Data Fig. 8c).

**Figure 4.**
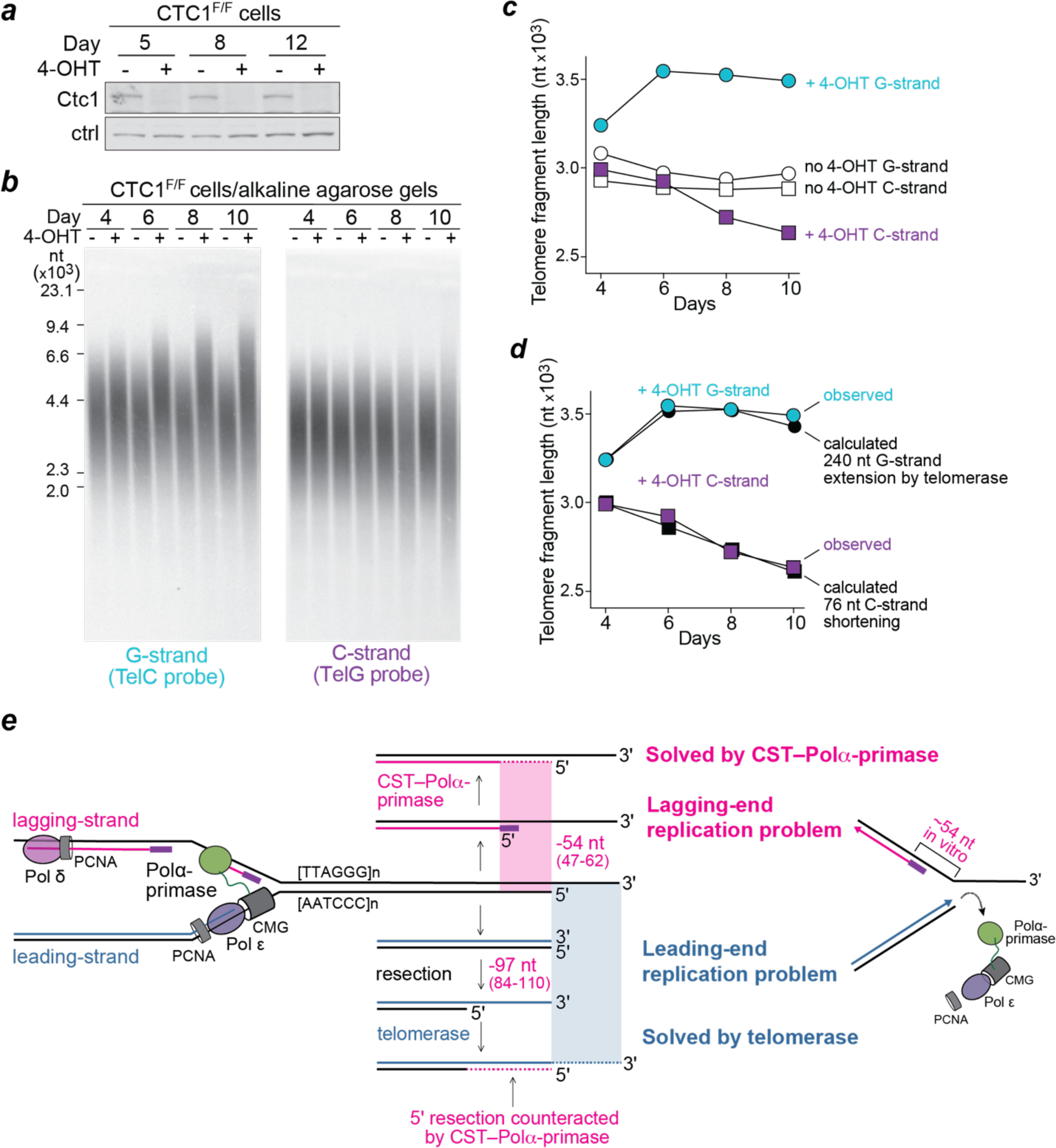
C-strand shortening at the predicted rate in absence of CST. ***a***, Immunoblot for Ctc1 in CTC1^F/F^ HCT116 cells with and without induction of Cre with 4-OHT. Ctrl: non-specific band used as a loading control. ***b***, Alkaline agarose gel analysis of the G- and C-rich of telomeric restriction fragments in telomerase proficient CTC1^F/F^ cells with or without 4-OHT treatment. Left: gel hybridized with TelC to detect the G-strands. Right, duplicate gel probed with TelG to detect the C-strands. ***c***, Graph showing the changes in length of the G- and C-strands as determined in (***b***). ***d***, Close agreement of the observed length changes in Ctc1-deficient cells with calculated length changes based on C-strand shortening of 76 nt/PD and G-strand elongation of 240 nt/PD by telomerase (see Extended Data Figure 9). ***e***, Schematic illustrating the two end-replication problems. The leading-end replication problem is solved by telomerase elongating the G-rich strand. CST–Polα-primase provides the solution to the lagging-strand replication problem which results from the position of the last Okazaki fragment synthesized by the replisome. In addition, CST–Polα-primase makes up for the C-strand sequence loss resulting from 5ʹ end resection.

The data reported here indicate that DNA replication creates a dual problem at telomere ends (Fig. 4e). The results of *in vitro* replication verify the long-held view that leading-strand replication leads to loss of the G-rich 3ʹ overhang sequence at one daughter telomere, representing the end-replication problem that can be solved by telomerase. We document a second end-replication problem *in vitro* and *in vivo* that results from the inability of lagging-strand DNA synthesis to synthesize a C-strand of the same length as the parental C-strand (Fig. 4e). Our data indicate that CST–Polα-primase counteracts this C-strand loss through fill-in synthesis at lagging-end telomeres. CST–Polα-primase-mediated fill-in also mitigates C-strand loss at the leading-end telomeres where resection generates the 3ʹ overhang used by telomerase^10,11,32,33^ (Fig. 4e). Together these processes shorten the telomeric C-strand by approximately 76 nt per population doubling in HCT116 cells. In keeping with the critical role of CST–Polα-primase in solving the second end-replication problem, defects in this pathway lead to telomere biology disorders –predominantly Coats plus and, more rarely, dyskeratosis congenita– (reviewed in refs. 21 and 40^21,40^). It was proposed that CST– Polα-primase evolved in LECA (or its Asgard archaeal ancestor) to maintain the 5ʹ ends of linear chromosomes^21,25^. Our data argue for this view since the lagging-strand end-replication problem likely threatened the maintenance of the first linear chromosomes.

## Methods

### Construction of replication templates

The 3ʹ-overhang and no-overhang templates used for *in vitro* DNA replication reactions were constructed by ligating oligonucleotides to the plasmid vVA25 following its linearization with SapI. To generate vVA25, a single BbvCI site was inserted ∼ 200 nt upstream of the SapI restriction site in vVA22^18^. For the 3ʹ-overhang template, a 25 bp duplex with a 100 base 3ʹ overhang was generated by annealing oligonucleotide VA67 (Sigma) and 3ʹ-overhang (IDT). To generate a blunt-ended template with the same sequence as the 3’ overhang, oligonucleotide VA67 was annealed to oligo JY556 (IDT). Annealing was performed by heating the oligonucleotides to 85°C for 5 min and then slow cooling to room temperature in 200 mM NaCl, 5 mM EDTA. Annealed oligonucleotides were then ligated onto SapI-linearized vVA25. For the 3ʹ overhang template, ligation (250 μl reaction) was performed overnight at 16°C in 1X T4 Ligase buffer with 50 nM SapI-linearized vVA25, 1.25 μM annealed oligonucleotides and 12,000 units T4 Ligase (NEB). For the no-overhang template, ligation (200 μl reaction) was performed overnight at 16°C in 1X T4 Ligase buffer supplemented with 5 mM Mg(OAc)_2_ and 0.5 mM ATP, 62.5 nM SapI-linearized vVA25, 1.56 μM annealed oligonucleotides and 12,000 units T4 Ligase (NEB). Ligation reactions were quenched by addition of EDTA to 25 mM and proteins were digested by incubation at 37°C for 20 min with 0.1% SDS and 20 units/ml proteinase K (NEB). Unligated oligonucleotides were removed by gel filtration through a 50 cm x 0.7 cm Sepharose 4B column (Sigma) equilibrated in 5 mM Tris-HCl (pH 8), 0.1 mM EDTA.

To construct in vitro DNA replication templates containing human telomeric repeats, eight TTAGGG repeats flanked by an AscI restriction site were inserted into vVA25 ∼ 250 bp downstream of the BbvCI site. The sequence of the resulting plasmid was verified by Sanger and Nanopore sequencing (Source Bioscience). Linear templates were prepared by digesting CsCl density gradient purified DNA with AscI (NEB) (terminal repeats template) or PciI (NEB) (internal repeats template). Digest were quenched by addition of EDTA to 25 mM and proteins were digested by incubation at 37°C for 40 min following addition of SDS and Proteinase K (NEB) to 0.25% and 20 units/ml respectively. Proteins were removed by phenol:chloroform:isoamyl alcohol (25:24:1) (PCI) (Sigma) extraction followed by ethanol precipitation. DNA was resuspended in 10 mM Tris-Cl (pH 8), 1 mM EDTA.

Oligos used for generation of DNA termini: VA67, 5ʹ-TAGTCCATCGGTTTTGCCATAAGAC-3ʹ; 3ʹ-overhang, 5ʹ-[Phos]GCTGTCTTATGGCAAAACCGATGGACTATGTTTCGGGTAGCACCAGGAGTCT GTAGCACGTGCATCTCAACGTGGCGTGAGTACCTTTTAATCACCGCTTCATGCTAAG GATCTGGCTGCATGCTATG-3ʹ; JY556 5ʹ-CATAGCATGCAGCCAGATCCTTAGCATGAAGCGGTGATTAAAAGGTACTCACGCCACGTTGAGATGCACGTGCTACAGACTCCTGGTGCTACCCGAAACATAGTCCATCGGTTTTGCCATAAGAC-3ʹ.

### Standard replication reaction

Standard replication reactions were carried out as reported previously^18^ with minor alterations. MCM loading and phosphorylation was performed in a reaction (55 μl) containing: 25 mM Hepes-KOH, pH 7.6; 100 mM potassium glutamate; 40 mM KCl; 0.01% NP-40-S; 1 mM DTT; 10 mM Mg(OAc)_2_; 0.1 mg/ml BSA; 3 mM ATP; 3 nM DNA template; 75 nM Cdt1-Mcm2-7; 40 nM Cdc6; 20 nM ORC; 25 mM DDK. After incubation at 24°C for 10 min, S-CDK was added to 80 nM and the reaction was incubated for a further 5 min at 24°C. The MCM loading reaction was then diluted 4-fold into replication buffer to give final replication reaction conditions of: 25 mM Hepes-KOH, pH 7.6; 257 mM potassium glutamate; 10 mM KCl; 0.01% NP-40-S; 1 mM DTT; 10 mM Mg(OAc)_2_; 0.1 mg/ml BSA; 3 mM ATP; 0.2 mM GTP; 0.2 mM CTP; 0.2 mM UTP; 30 μM dATP; 30 μM dTTP; 30 μM dCTP; 30 μM dGTP; 0.75 nM DNA template; 18.75 nM Cdt1-Mcm2-7, 10 nM Cdc6; 5 nM ORC; 6.25 nM DDK; 20 nM S-CDK and either 33 nM α-[^32^P]-dCTP (Hartmann Analytic #SCP-205) (Fig. 1d) or 33 nM α-[^32^P]-dATP (Hartmann Analytic #SCP-203) (Fig. 1f and Extended Data Figs. 2e and 3b). Reactions (100 μl) were equilibrated at 30°C and DNA replication was initiated by addition of replication proteins from a master mix to final concentrations of: 30 nM Dpb11; 100 nM GINS; 30 nM Cdc45; 10 nM Mcm10; 15 nM Pol ε; 20 nM Ctf4; 100 nM RPA; 20 nM RFC; 20 nM Tof1-Csm3; 20 nM PCNA; 5 nM Pol δ; 12.5 nM Sld3-7; 20 nM Sld2; 10 nM Mrc1; 20 nM Fen1; 20 nM Ligase and, unless indicated otherwise, 20 nM Polα-primase. Protein storage buffers additionally contributed approximately 12 mM NaCl/KCl and 25 mM KOAc/K-glutamate to the reactions. Reactions were incubated at 30°C for 30 min and were quenched by addition of EDTA to 50 mM. Proteins were removed by treatment with proteinase K (20 units/ml) (NEB) and SDS (0.25%) for 20 min at 37°C followed by PCI extraction. Unincorporated nucleotide was removed using Illustra G-50 columns (GE Healthcare).

To map the 5ʹ and 3ʹ ends generated by the rightward moving replication fork when it runs off the end of the template, products were first digested (or mock digested) with RNase HII. Deproteinized reactions (63 μl) were incubated for 45 min at 37°C in 1X ThermoPol buffer (NEB) with 15 units *E. coli* RNase HII (NEB #M0288). Reactions were quenched by addition of EDTA to 50 mM and proteins were removed by treatment with proteinase K (20 units/ml) (NEB) and SDS (0.25%) for 10 min at 37°C followed by PCI (Sigma) extraction. The DNA was ethanol precipitated following addition of 40 μg glycogen (Invitrogen) and was resuspended in 65 μl 1X rCutSmart buffer (NEB). Samples were divided into three 20 μl aliquots that were treated (or mock treated) for 30 min at 37°C with 10 units of Nt.BbvCI or Nb.BbvCI as indicated in the figures. Reactions were quenched by addition of EDTA to 50 mM and proteins were removed by treatment with proteinase K (20 units/ml) (NEB) and SDS (0.25%) for 10 min at 37°C followed by PCI extraction and ethanol precipitation. For analysis through denaturing urea polyacrylamide gels, DNA was resuspended in 4 μl 10 mM Tris-Cl (pH 8), 1 mM EDTA and then supplemented with an equal volume of 2X loading buffer (95% formamide, 20 mM EDTA, 0.04% xylene cyanol, 0.04% bromophenol blue). After incubation at 95°C for 5 min, products were immediately resolved on 5 - 5.5%, 40 cm x 20 cm polyacrylamide (Bis-Acrylamide 19:1 – Fisher Scientific), 7 M Urea gels, run in 1X Tris-Borate-EDTA buffer (TBE) for 130 min at 40 W. Gels were dried under vacuum at 75°C onto 3MM paper (Whatman) and imaged using Typhoon phosphorimager (GE Healthcare).

### Analysis of in vitro DNA replication reactions

To generate normalized plots of DNA replication reactions, lane profiles were generated in ImageJ for each reaction condition and the molecular weight markers (50 bp ladder, NEB, #N3236). Background was subtracted from each lane profile by subtracting the signal from the equivalent reaction condition (-/+ RNase HII) but without BbvCI digestion. Following background subtraction, data were normalized by dividing the pixel intensity at each migration position by the pixel intensity of the most intense product band within the region displayed in each plot. To determine the approximate length of replication products in nucleotides, the relative migration distance (in centimeters) of each marker band was plotted against the log10 of its length (in nucleotides). Data were fit to a third order polynomial using Prism (GraphPad) and the resulting fits were used to interpolate product length values (in nucleotides) for every migration position of a given lane profile.

### Cell culture, deletion of CTC1, depletion of telomerase, and immunoblots

CTC1^F/F^ HCT116 cells were cultured as described^32^. For *CTC1* deletion, the cells were treated with 0.5 μM 4-hydroxytamoxifen (4-OHT) for 5 h in media to induce Cre recombinase. Ctc1 loss was verified by immunoblotting as described ^33^ using rabbit polyclonal antibody raised against full length recombinant human Ctc1 (a gift from John C. Zinder). CTC1^F/F^ cells were rendered telomerase deficient by bulk targeting of hTERT with lentiCRISPR v2 vector expressing Cas9 and sgRNAs (5ʹ-ACACGGTGACCGACGCACTG-3ʹ and 5ʹ-CTTGGTCTCGGCGTACACCG-3ʹ). Telomerase was further inhibited using 16 μM BIBR1532^35^ (TOCRIS #2981) in the media. TRAP assays for telomerase activity were performed using the TRAPeze kit (Millipore, #S7700) according to the manufacture’s instruction.

### Overhang analysis of leading- and lagging-end telomeres

Isolation of genomic DNA and separation of leading- and lagging-end telomeres were performed as described^10,14^. Briefly, cells were cultured with 100 μM BrdU for 18 h, harvested, and genomic DNA was extracted. DNAs were digested with *Mbo*I/*Alu*I and fractionated on CsCl density gradients and fractions were collected from the bottom of the centrifuge tubes. Telomeric DNA in each fraction was quantified by slot blot hybridization using γ-^32^P-ATP end-labeled [TTAGGG]_4_ (TelG probe) as described^10^. Leading- and lagging-strand peak fractions were pooled, dialyzed to remove most CsCl, and DNAs were isolated by ethanol precipitation followed by suspension in TE. The pooled fractions were analyzed using the in-gel overhang assay with a γ-^32^P-ATP end-labeled [AACCCT]_4_ (TelC probe) as described^10^.

### PCR assay for telomeric overhang (Method I)

2×10^6^ cells were trypsinized, pelleted, frozen in liquid N_2,_ and stored at -80°C. For DNA extraction, the cell pellet was resuspended to 1 ml TLE buffer containing 10 mM Tris-HCl pH 7.8, 100 mM LiCl, 10 mM EDTA, and 100 μg/ml RNase A (Sigma, #R5000). 1 ml TLES buffer (TLE + 0.1% (w/v) SDS) with 100 μg/ml proteinase K (Roche, #03115879001) was added and the suspension was incubated at 37°C for 10 min. Genomic DNA was extracted with 2 ml phenol-chloroform-isoamyl alcohol (PCI, Thermo Fisher, #BP1752I), followed by DNA precipitation with 1/20 volume of 8 M LiCl and 1 volume of isopropanol. Genomic DNA was resuspended in 50 μl TE buffer containing 50 mM LiCl (50 mM LiCl-TE), 100 μg/ml RNase A, and 0.2 U/μl RNase H (NEB #M0297) and immediately used for the overhang assay as follows. Genomic DNA (30 μl) was poly(A) tailed using 20 units terminal transferase (TdT, NEB, #M0315) and 0.2 mM dATP for 15 min at 37°C in TdT buffer (20 mM Tris-HCl pH 7.9, 50 mM LiCl, 10 mM MgCl_2_), followed by PCI extraction and isopropanol precipitation as above. DNA was resuspended in 40 μl Extension buffer (10 mM Tris-HCl pH 7.9, 50 mM LiCl, 10 mM MgCl_2_, and 50 μg/ml BSA). The tailed DNA was annealed to 0.2 nM 5ʹ biotinylated teltail ss-adaptor at 21°C for 10 min. Primer extension was performed with 0.006 U/μl Q5 (NEB, #M0491) and 200 μM each of dATP/dCTP/dTTP at 21°C for 10 min and an additional 15 min incubation with 25 nM recombinant human RPA (a gift from Sarah W. Cai) and 0.03 U/μl T4 DNA polymerase (NEB, #M0203) at 37°C. The teltail ss-adaptor (see below) is a mixture of single-strand DNA oligos of 5ʹ biotinylated teltail^37^ followed by [T]_8_ and 4 nt of C-strand sequence in 6 permutations. Extended DNA was digested with *Alu*I at 37°C for 20 min to diminish the viscosity. Excess teltail ss-adaptor and short bulk genomic DNA fragments were removed by DNA size selection using 25 μl SPRI select (Beckman, 0.5x v/v SPRI beads to DNA suspension). After 2 washes with 80% EtOH according to the manufacturer’s instructions, DNA was eluted in 60 μl 50 mM LiCl-TE buffer and the concentration of DNA was measured using NanoDrop (Thermo Fisher). 0.5 μg DNA was used for the subsequent steps. After capture of the products on washed Streptavidin beads (10 μl, Dynabeads M-280; Thermo Fisher, #11205D), the DNA was denatured with 150 mM NaOH for 5 min at 21°C 2 times, neutralized with 2 washes with 1 M LiCl-TE buffer, one wash with 50 mM LiCl-TE, and resuspended in 50 μl TdT buffer containing 2 mM rGTP (Promega, E603B) and 20 U TdT at 37°C for 15 min for rGTP tailing^41^. DNA-bound beads were rinsed 3 times with 50 mM LiCl-TE and ligated to P1 ds-adaptor for >1 h at 21°C in Ligation mix (Extension buffer containing 20 nM P1 ds-adaptor, 200 μM ATP, 25% PEG 8000, 8 U/μl Salt-T4 DNA ligase (NEB #M0467)). P1 ds-adaptors are double-strand (ds) splint adaptors that contain 5ʹ overhangs with the 6 permutations of the telomeric sequence (see below). 6 permutated upper-oligos and the 5ʹ phosphorylated bottom strand oligo were mixed, annealed, and kept as aliquots at -20°C to avoid multiple freeze-thaw cycles. After 2 washes with 1 M LiCl-TE buffer, DNA-bound beads were resuspended in 20 μl TE buffer.

PCR with the teltail and P1 PCR primers was performed using Q5 PCR enzyme mix (NEB, #M0494) and the following conditions using 0.1 μl DNA-bound beads: denaturation at 98°C for 30 sec; 30 cycles of 10 sec at 98°C, 20 sec at 69°C, and 50 sec at 72°C; final extension for 5 min at 72°C. PCR products were denatured at 80°C for 5 min in 2x loading buffer containing 95% Formaldehyde, 10 mM EDTA, 0.05% xylene cyanol (Sigma, X4126), and 0.05% bromophenol blue (Sigma B-1256) and separated on 4.5% polyacrylamide/7 M urea gels in 1xTBE buffer and electroblotted in 0.5xTBE onto nylon membranes (Hybond-N^+^, Cytiva). DNA was UV-crosslinked to the membranes, hybridized with γ-^32^P-ATP end-labeled [TTAGGG]_4_ (TelG probe) in Church mix at 48°C for overnight and washed three times in 4x SSC and 1 time in 4x SSC/0.1% SDS at 48°C for 5 min each wash. Imaging was done with a Typhoon phosphorimager.

For PCR assay with synthetic telomeric oligos [TTAGGG]_5-20_-TTAG-[A]_10_, 20 pM ss-oligo DNA was used for primer extension. The poly(A) tailing, *Alu*I digestion, and DNA size selection steps were skipped. The standard curve was generated using GraphPad Prism software (Dotmatics, ver.10).

### Generation of the model telomere substrate

A model telomere substrate containing a 56 nt 3’ telomeric overhang was constructed using the telomeric DNA containing *Bgl*II/*Hind*III fragment from pSXneo1.6T_2_AG_3_^42^. To generate the 3ʹ overhang, 1 μM 5ʹ-CTAACCCTAAGCTCTGCGACAT -3ʹ and 5ʹ-GATCATGTCGCAGAGC[TTAGGG]_11_ -3ʹ were phosphorylated with T4 polynucleotide kinase, annealed, and incubated with 0.1 μM linear pSXneo1.6T_2_AG_3_. Ligation was done in the presence of *Bgl*II and *Hind*III with 6 cycles at 37°C for 5 min and 21°C for 5 min. The ligated product was gel purified on a 1% agarose gel.

### *Exo*I digestion

Genomic DNA and the model telomere substrate were incubated with 40 U *E. coli Exo*I (#M0293, NEB) in 67 mM Glycine-NaOH (pH 9.5), 6.7 mM MgCl_2_ and 10 mM 2-mercaptoethanol at 37°C for >6 h. After *Exo*I digestion, DNA was purified by PCI extraction and precipitated by adding 1 volume of isopropanol and 1/20 volume of 8 M LiCl. For in-gel and Southern blot telomere hybridization assays, genomic DNA was digested with *Mbo*I and *Alu*I after the *Exo*I treatment.

### Oligonucleotides for the PCR method

Oligonucleotides were synthesized by IDT or Thermo Fisher with indicated modifications and purification on PAGE.

The teltail ss adaptors 1-6 used for extension were as follows (telomeric sequences underlined). Adaptor 1: 5ʹ-[5biotin]-TGCTCCGTGCATCTGGCATC TTTTTTTT <u>CTAA</u>-3ʹ; adaptor 2: as adaptor 1 but ending in <u>TAAC</u>-3ʹ; adaptor 3: as adaptor 1 but ending in <u>AACC</u>-3ʹ; adaptor 4: as adaptor 1 but ending in <u>ACCC</u>-3ʹ; adaptor 5: as adaptor 1 but ending <u>CCCT</u>-3ʹ; adaptor 6: as adaptor 1 but ending in <u>CCTA</u>-3ʹ.

The P1 ds adaptors upper 1-6 oligonucleotides were as follows (telomeric sequences underlined). Upper 1: 5ʹ-[5AmMC6]-GCACAGCCACTGGTAACG CCC <u>GGTTAG</u>-[3AmC7]-3ʹ; upper 2, as upper 1 but ending in <u>GTTAGG</u>-[3AmC7]-3ʹ; upper 3: as adaptor 1 but ending in <u>TTAGGG</u>-[3AmC7]-3ʹ; upper 4: as adaptor 1 but ending in <u>TAGGGT</u>-[3AmC7]-3ʹ; upper 5: as adaptor 1 but ending in <u>AGGGTT</u>-[3AmC7]-3ʹ; upper 6: as adaptor 1 but ending in <u>GGGTTA</u>-[3AmC7]-3ʹ. The P1 ds adaptor lower was: 5ʹ-[5phos]-CGTTACCAGTGGCTGTGC-[3ddC]-3ʹ.

PCR was performed with the following primers. Teltail primer: 5ʹ-TGCTCCGTGCATCTGGCATC-3ʹ; P1 primer: 5ʹ-GCACAGCCACTGGTAACG-3ʹ The following control oligos were used to test the assay. Tel5-TTAG-A10: 5ʹ-[TTAGGG]_5_TTAG AAAAAAAAAA-3ʹ; Tel10-TTAG-A10: 5ʹ-[TTAGGG]_10_TTAG AAAAAAAAAA-3ʹ; Tel15-TTAG-A10: 5ʹ-[TTAGGG]_15_TTAG AAAAAAAAAA-3ʹ; Tel20-TTAG-A10: 5’-[TTAGGG]_20_TTAG AAAAAAAAAA-3ʹ.

### Telomeric overhang measurement (Method II)

In method II, genomic DNA isolated as described previously^43^ was treated (or not) with *E. coli Exo*I as described above, digested with *Mbo*I and *Alu*I, PCI extracted, isopropanol precipitated, and resuspended in 2 mM Tris-HCl (pH 7.4) containing 0.2 mM EDTA. 1.5 μg DNA was suspended in Alkaline loading buffer containing 50 mM NaOH, 1 mM EDTA, 3% Ficoll 400 (Sigma, F9378), 0.5% (w/v) bromocresol green, 0.9% (w/v) xylene cyanol FF, fractionated on alkaline gels (0.7% agar, 50 mM NaOH and 1mM EDTA) in alkaline running buffer contains 50 mM NaOH and 1 mM EDTA as described^43^. The gels were neutralized in neutralizing buffer (0.5 M Tris-HCl pH 7.0 and 3 M NaCl), dried and processed for in-gel overhang assays with end-labeled TelC as described above. Gels were scanned using a Typhoon phosphorimager, and the hybridization intensity of the scan was read and analyzed using Fiji software (ImageJ version 2.0). The average size of the fragments was calculated taking into account that the hybridization intensity increases with fragment length^44^. To account for the subtelomeric component of the TRFs (0.5 kb) the mean TRF (Telomere Restriction Fragment) length was calculated as: Mean TRF = Σ(SI*i*)/Σ{SI*i* / (MW*i*-0.5)}+0.5, where SI is the signal intensity at position *I* and MW*i* is the TRF length at that position. Overhang lengths were calculated by comparing the G-strand length before and after *Exo*I treatment.

As a control, pSXNeo1.6T_2_AG_3_^42^ was digested with *Hind*III and *Bgl*II, ligated (or not) to an oligonucleotide bearing a telomeric 3ʹ overhang of 56 nt (as described above for the model telomere substrate), and the 3 kb fragments with and without the overhang were purified from agarose gel. The DNA fragments were treated (or not) with 20 U of *E. coli Exo*I, processed on alkaline gels as above, and detected by in-gel hybridization with the TelC probe.

### DNA markers for telomere blotting

For telomere markers, pSXneo1.6T_2_AG_3_ was digested by following combinations of restriction enzymes (NEB), and fragments of the corresponding sizes were purified from 1% agarose TAE gel electrophoresis. 5366 bp marker: *Bgl*II in NEB buffer r3.1; 4470 bp: *Pvu*II/*Bgl*II (r3.1); 2965 bp: HindIII/*Bgl*II (r3.1); 887 bp: *Kpn*I-HF in rCutSmart (rCS); 3304 bp: *Pci*I/*Nde*I (r3.1); 3121 bp: *Hinc*II/*Nde*I (rCS); 2814 bp: *Nae*I/*Nde*I (rCS); 2616 bp: *Bsa*I/*Nde*I/*Sca*I-HF (rCS); 3213 bp: *Pci*I/*Bgl*II (r3.1); 3017 bp: *Hinc*II/*Not*I-HF (rCS); 2710 bp: NaeI/*Not*I-HF/*Sca*I-HF (rCS); 2512 bp: *Bsa*AI/*Not*I-HF (rCS). 10 pg DNA of each marker was loaded per lane for gel electrophoresis. Each fragment contains 1.5 kb telomeric repeat sequences, with the exception of the 887 bp *Kpn*I-HF fragment which contains 0.8 kb telomeric repeats. The markers were detected by hybridization to the TelC probe. For the detection of 1 kb DNA ladder (NEB, #N3232) and 𝜆 *Hind*III/𝜑X *Hae*III marker, DNA ladders were denatured, end-labeled using γ-^32^P-ATP, and used as probes. Standard curves of the DNA markers were generated using GraphPad Prizm software (version 10.0) and the equation of the standard curve was generated using https://mycurvefit.com.

### Alkaline gel analysis of the telomeric G- and C-strands

Genomic DNA from was digested with *Mbo*I and *Alu*I and denatured in 50 mM NaOH and 1 mM EDTA before fractionating the DNA on 0.7% alkaline agarose gels as described above. Gels were neutralized in 0.5 M Tris-HCl (pH 7.0)/3 M NaCl, dried, prehybridized in Church mix for 1 h at 50°C, and hybridized overnight at 50°C in Church mix with γ-^32^P-ATP end-labeled [AACCCT]_4_ (TelC) or [TTAGGG]_4_ (TelG) as described^10^.

### Statistics and reproducibility

The experiments in Fig.1d and e and Extended Data Figs. 2e and 3e were performed a minimum of 3 times. For all other experiments, the number of replicates and statistics are indicated in the figures and figure legends.

## Author contribution

H.T. and T.d.L. conceived and designed the study. A.V. and J.T.P.Y. designed and performed the *in vitro* replication experiments. All *in vivo* experiments were executed by H.T. with help from P.B. in the early stages of the project. T.d.L. and J.T.P.Y. wrote the manuscript with input from all authors.

## Acknowledgements

We thank Carolyn Price for providing the CTC1^F/F^ HCT116 cell line. We thank the members of the de Lange lab for comments on this work, Viviana Risca and Andres Mansisidor for advice on the PCR method, John C. Zinder for the Ctc1 antibody, and Sarah Cai for purified human RPA. This work is supported by grants from the NCI (R35CA210036), the NIA (RO1AG016642), and the BCRF to TdL; a grant from the NCI to HT (5R50CA243771-02); and a grant to JTPY from the Medical Research Council (No. MC_UP_1201/12).

## Extended Data Figure Legends

**Extended Data Figure 1.**
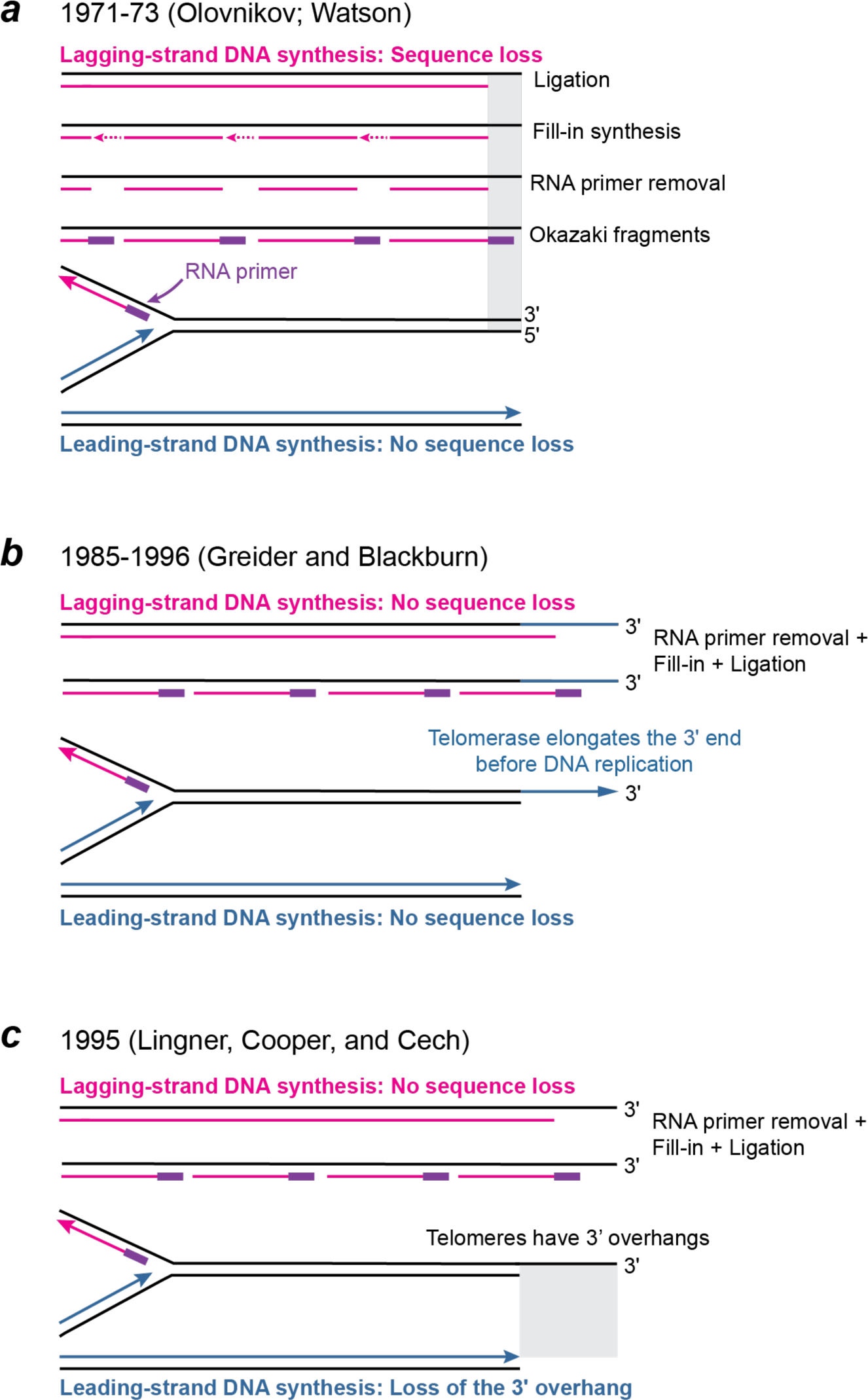
Changing views of the end-replication problem. *a*, The first version of the end-replication problem as perceived by Watson^5^ and Olovnikov^6^ after it was shown that Okazaki fragments carry RNA primers at their 5’ ends. In this end-replication problem, the 5’ ended strand of the lagging-strand DNA synthesis product is shorter than the parental strand. *b*, Telomerase was suggested to solve the end-replication problem by extending the 3’ ends before DNA replication^7^. *c*, When it had become clear that telomeres ended in a 3’ overhang, it was argued that the end-replication problem involved the inability of leading-strand DNA synthesis to recreate this part of the telomeric DNA^2^. Therefore, the end-replication problem involved the shortening of the G-rich strand at leading-end telomeres. Telomerase can solve the leading-end replication problem by acting after DNA replication (and 5’ end resection) as shown in Fig. 1a. If the last Okazaki fragments start on the 3’ overhang (as shown), no sequence loss will occur at lagging-end telomeres. The end-replication problem was therefore “no longer a lagging strand problem” (quote from ref. 2 ^2^).

**Extended Data Figure 2.**
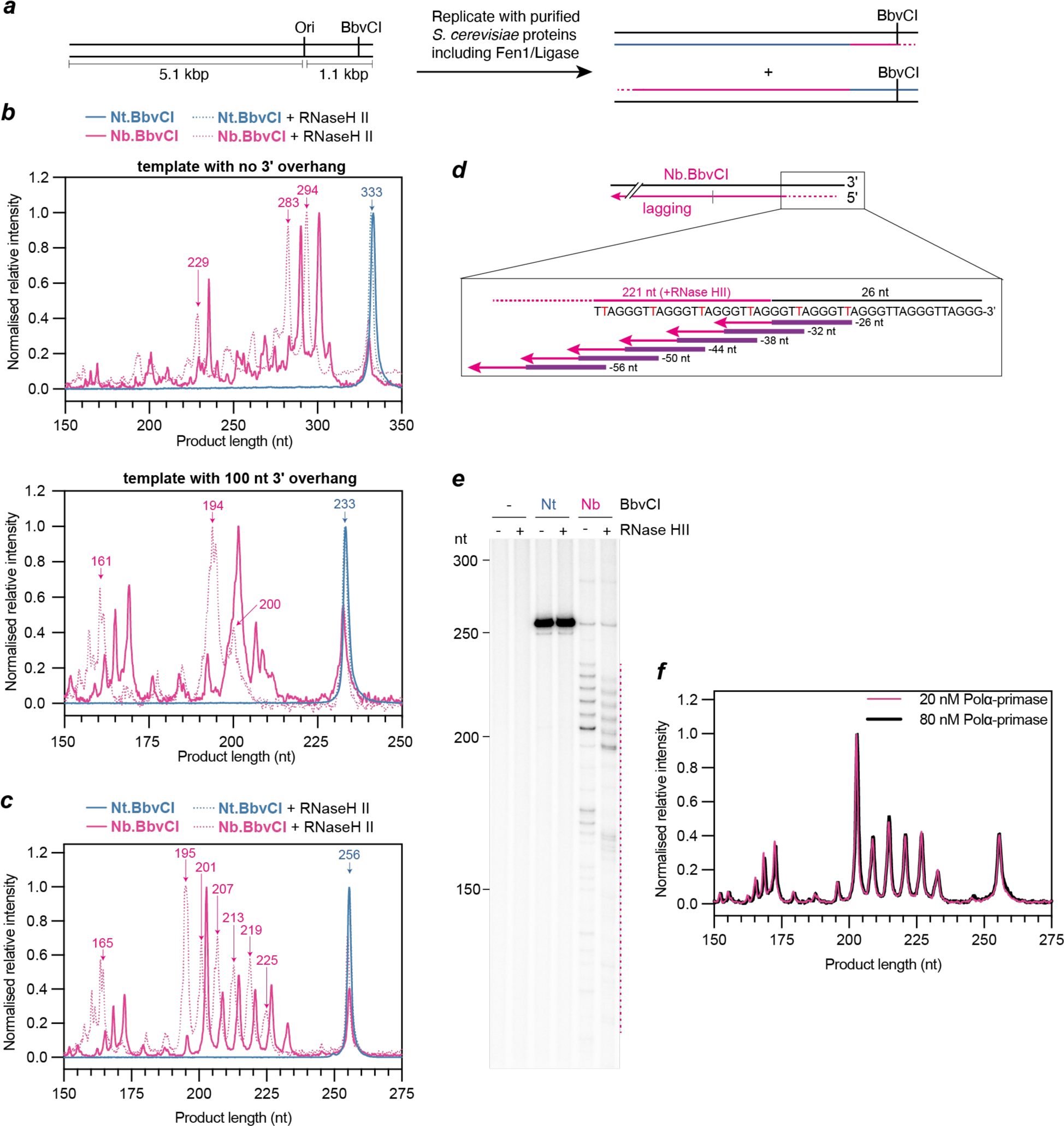
Analysis of replisome-mediated DNA synthesis at the end of linear DNA templates. ***a***, Schematic of the replication templates used for *in vitro* budding yeast DNA replication and the anticipated reaction products of origin-dependent replication. **b** and **c**, Analysis of the data presented in Fig. 1d (**b**) and 1f (**d**). Product length was determined using a standard curve derived from the molecular weight standards. Replication product intensity was normalised by dividing each data point by the amplitude of the most intense product for a given condition. **d**, Schematic illustrating the terminal TTAGGG repeats that are used to initiate Okazaki fragment synthesis during in vitro DNA replication. **e**, Replication reaction performed and analyzed as in Fig. 1f but with the concentration of Polα-primase increased from 20 nM to 80 nM. To enable clear visualization of the Nb.BbvCI products, the Nt.BbvCI product bands are saturated**. f**. Comparison of Nb.BbvCI digested replication products in the absence of RNase HII at 20 nM (Fig1.d) and 80 nM (**a**) Polα-primase. Data were processed as described for **b** and **c**.

**Extended Data Figure 3.**
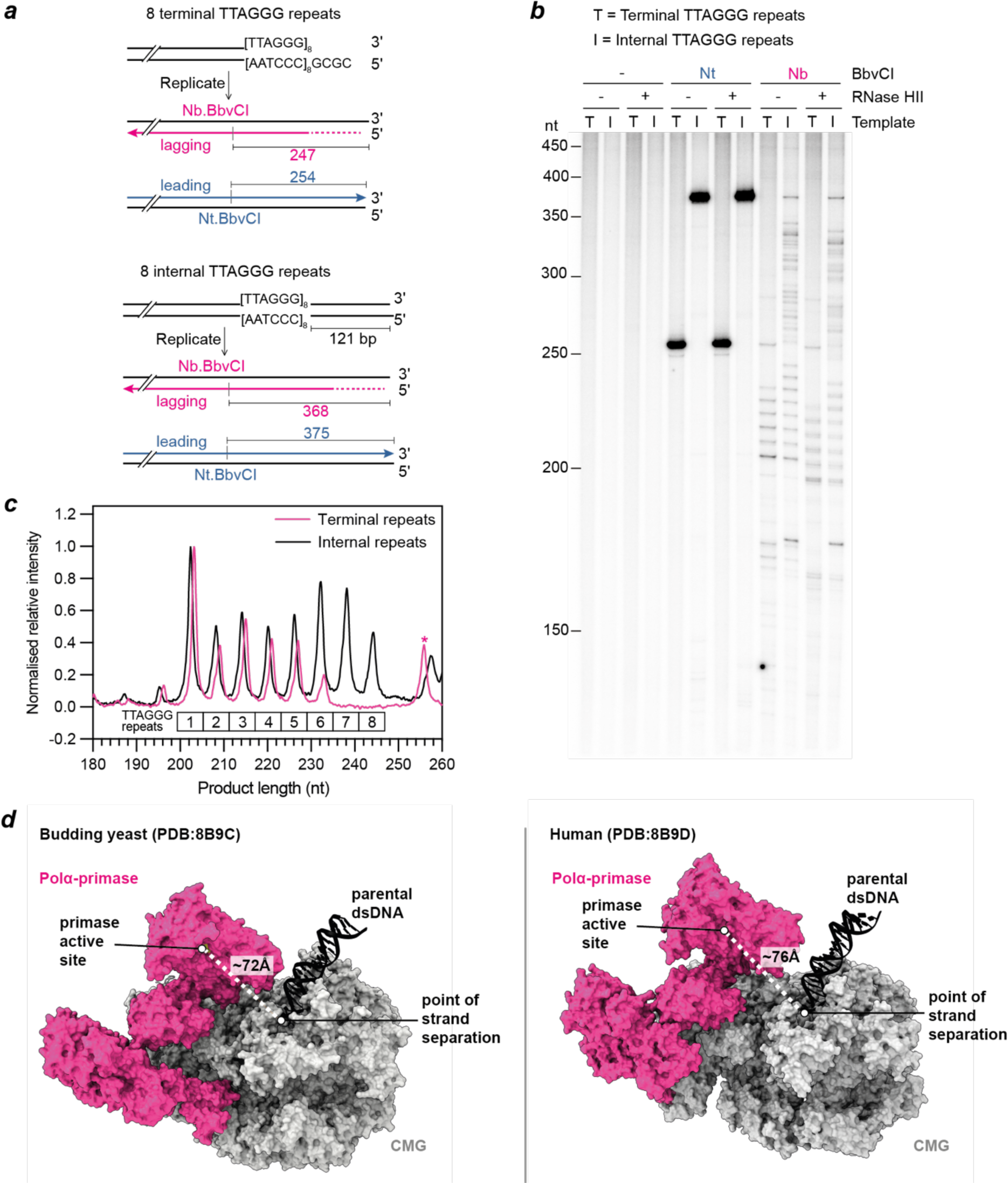
Analysis of lagging-strand priming within TTAGGG repeats. **a**. Schematic illustrating the TTAGGG repeat containing templates and the anticipated products of Nt./Nb.BbvCI digestion. By linearizing the plasmids with different restriction enzymes, the TTAGGG repeats are located either at the end of the template, or 121 bp from the end of the template. **b**. Denaturing 5% polyacrylamide/urea gel analysis of a replication reaction on the templates shown in (**d**) analyzed as described in Fig. 1d. To enable clear visualization of the Nb.BbvCI products, the Nt.BbvCI product bands are saturated**. c**. Comparison of Nb.BbvCI digested replication products in the absence of RNase HII from **b**. The asterix marks the presence of an RNase HII insensitive replication product. The position of the template strand TTAGGG repeats are illustrated below the traces. **d**, Comparison of Polα-primase in the yeast and human replisomes. For clarity only Polα-primase (pink), the CMG helicase (grey) and DNA (black) are shown. The shortest distance between the point of strand separation and the primase active site are illustrated (dashed white lines).

**Extended Data Figure 4.**
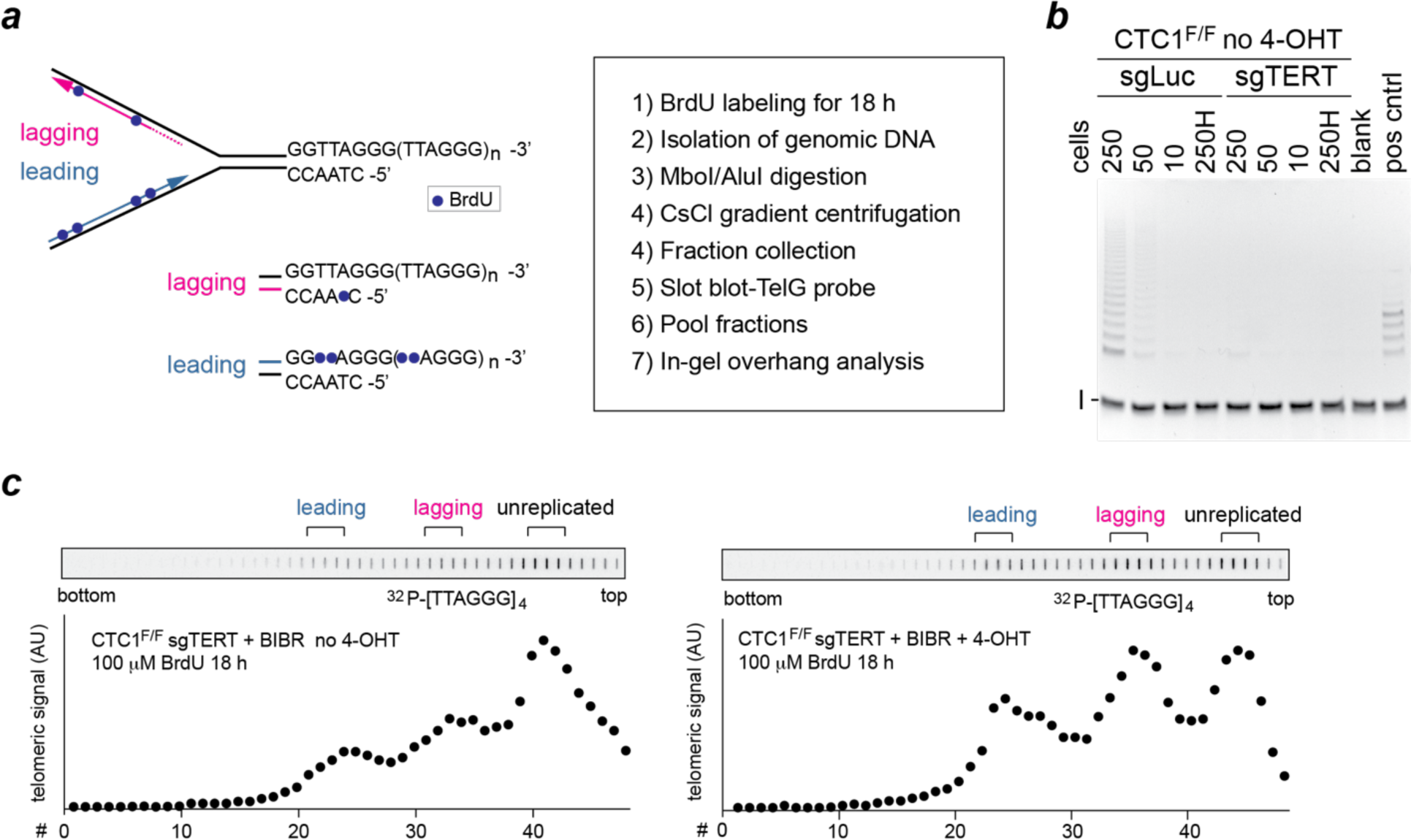
CsCl-gradient separation of leading- and lagging-end telomeres from CTC1^F/F^ cells lacking telomerase activity. ***a***, Schematic of the experimental approach to separate telomeres replicated by leading- and lagging-strand DNA synthesis. ***b***, TRAP assay on extracts from CTC1^F/F^ cells from which hTERT is deleted (in bulk) using CRISPR/Cas9 (sgTERT). Cells treated with a Luciferase sgRNA are used as the control. Cell equivalents are indicated above the lanes. 250H: Heat-inactivated extract (250 cells). Positive control sample (pos cntrl) provided by the manufacturer. I: internal PCR control. ***c***, Examples of slot blots to detect telomeric DNA in fractions from CsCl gradients. Pooled fractions used for in-gel 3ʹ overhang assays are indicated at the top.

**Extended Data Figure 5.**
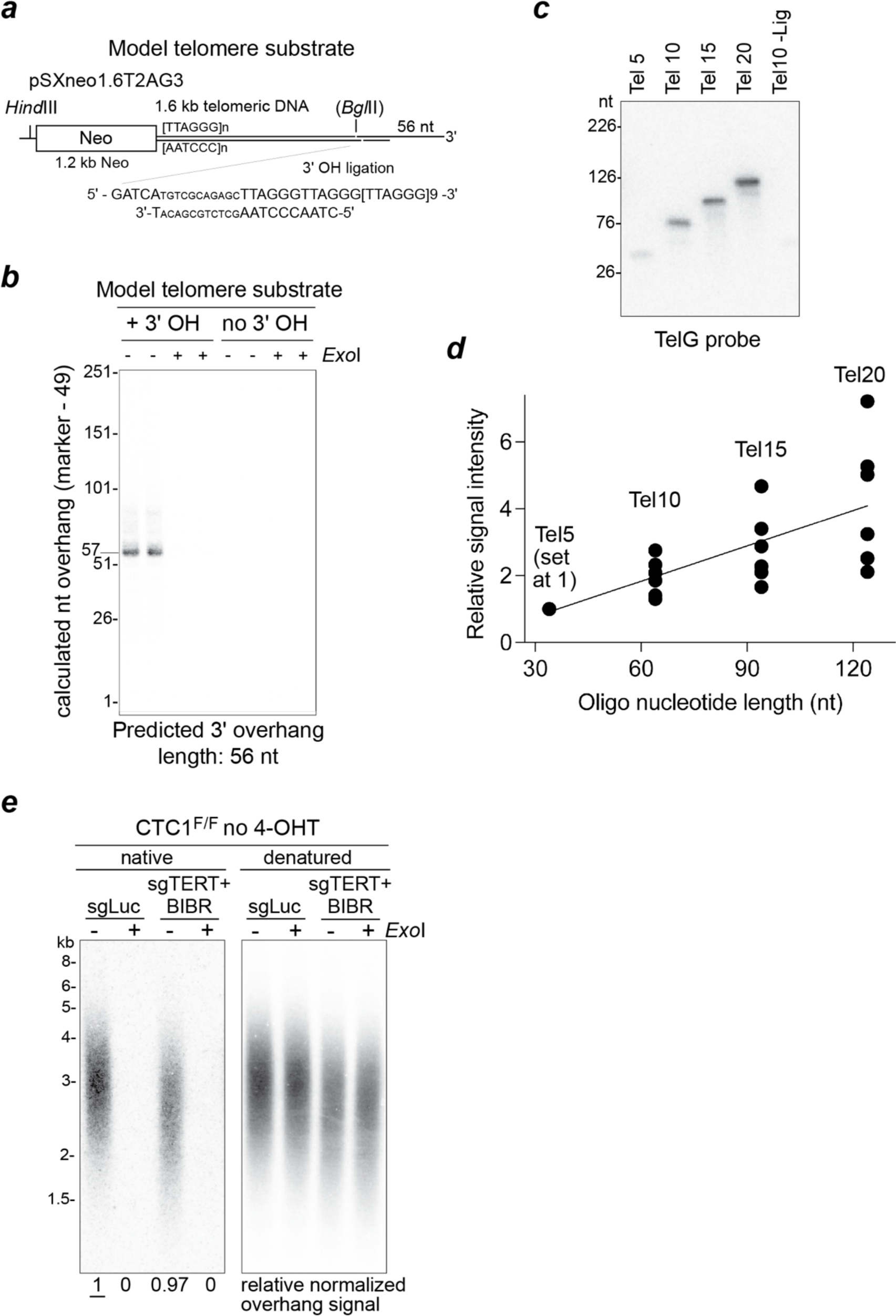
Validation of the PCR overhang assay (Method I) ***a***, Schematic of the telomere model substrate generated by ligating a telomeric 3ʹ overhang to a *Hind*III/*Bgl*II fragment of pSXneo1.6T_2_AG_3_. ***b***, PCR overhang assay on the model telomeric substrate before and after *E. coli* 3’ exonuclease *Exo*I treatment to remove the 3’ overhang. ***c***, PCR overhang assays performed with synthetic telomeric oligos containing 5 to 20 TTAGGG repeats. Tel10 -Lig: Tel10 oligo processed in parallel but without ligation to the adaptor. ***d***, Relative signal intensity of 6 independent PCR overhang assays as in (***c***) were plotted. The signal intensity of Tel5 oligo in each experiment was set at 1. ***e***, In-gel hybridization assay showing that telomerase expression does not strongly alter 3ʹ overhang lengths when Ctc1-profient cells and that the *Exo*I digestion used in Fig. 2e worked. Genomic DNAs from CTC1^F/F^ cells with or without sgTERT and BIBR1532 treatment were treated with *E. coli Exo*I as indicated and digested with *Mbo*I/*Alu*I for the in-gel overhang assay shown. Left: TelC probe detecting the overhang signals. Right: same gel probed with TelC after in situ denaturation to detect the total telomeric DNA for normalization.

**Extended Data Figure 6.**
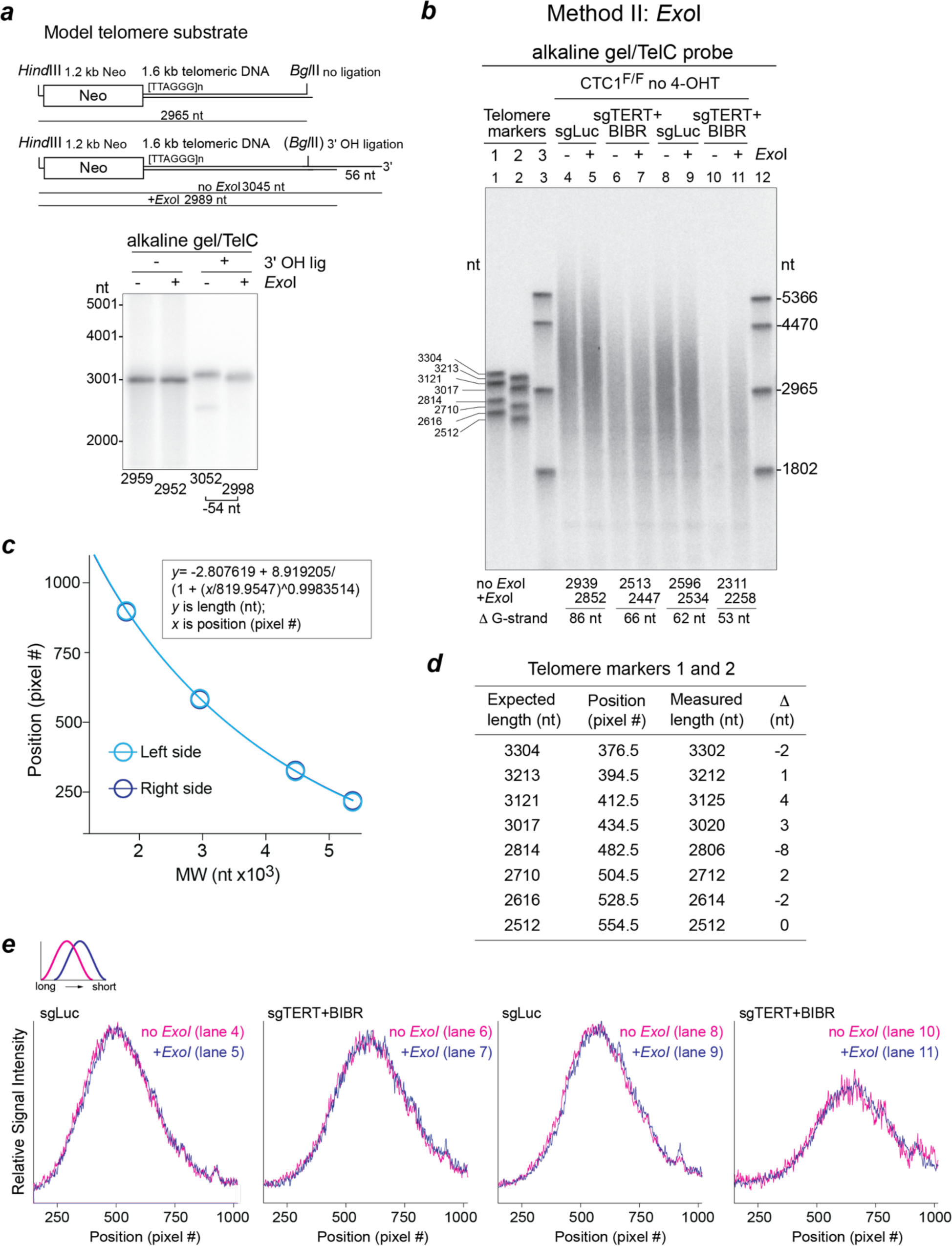
Overhang measurement with *Exo*I digestion (Method II) ***a***, Top: Telomere model substrate with or without overhang (as in Ext. Data Fig. 5a) with predicted fragment lengths indicated. Bottom: Telomere model substrate with and without overhang was treated with or without *Exo*I, separated by alkaline agarose gel electrophoresis and detected by in-gel hybridization with the TelC probe. Sizes of the detected fragments were calculated based on the molecular weight marker. Shortening of the model telomere substrate G-strand by *Exo*I is shown below the gel. ***b***, Example of determination of G-strand shortening by *Exo*I treatment. DNAs from cells with or without depletion of telomerase (sghTERT+BIBR) were treated with *Exo*I as indicated and then digested with *Mbo*I and *Alu*I. DNAs were separated on an alkaline agarose gel and the gel was hybridized with TelC. Changes in G-strand length after *Exo*I treatment are indicated below the lanes. The MW markers are generated by restriction digestion of pSXneoT2AG31.6 (see Extended Data Fig. 5a and methods) and fragments containing TTAGGG repeats are detected by TelC. ***c***, Graph of migration (pixel position) versus the MW of the telomeric marker in lane 3 and lane 12 of the gel (***b***) generated by scanning of the gel using Fiji software (ImageJ, ver2.0). The equation of the curve is used to convert migration (pixel #) to telomere lengths. ***d***, Measured length of the telomere markers in lane 1 and 2 was calculated using the position of the fragment (pixel #) in the gel and the equation generated from standard curve in (***c***) and compared to the expected length. ***e***, Overlay of the scan profiles of G-strand telomeric signal from DNA treated with or without *E. coli Exo*I shown in (***b***). Smaller pixel # represents longer telomere length. Line graphs shift to the right (lower MW) in DNAs treated with *Exo*I compared to mock-treated DNAs.

**Extended Data Figure 7.**
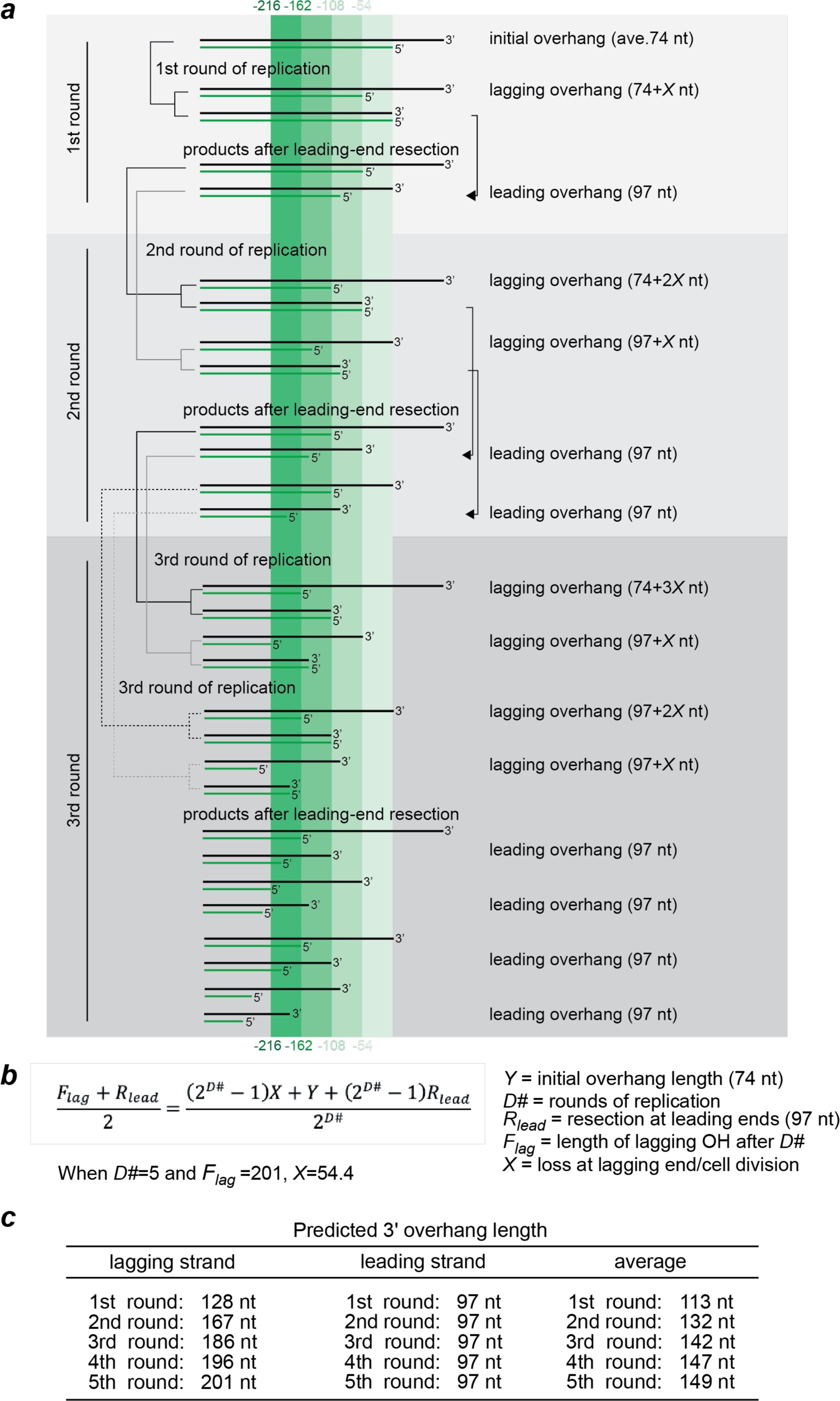
Calculation of C-strand shortening at lagging-end telomeres. ***a*** and ***b***, Calculation of the C-strand loss at lagging-end telomeres in cells lacking Ctc1 and telomerase activity. The schematic in (***a***) shows the changes at leading- and lagging-end telomeres over three rounds of assuming 97 nt (average of 110 and 84 nt) resection of the leading-end products as determined in Fig. 2j. Sequence loss at the lagging-end C-strand is calculated to be 54 nt based on the 3’ overhang length of 201 nt (average of 228 and 173) for lagging-end telomeres after 5 divisions without Ctc1 (Fig. 2 ***j***) and applying the equation in (***b)***. Note that the 97 nt loss at the leading-end telomeres is constant, whereas the loss at the lagging-end telomeres increases. ***c***, Predicted overhang length changes in 5 successive rounds of replication after CTC1 deletion from telomerase-deficient cells as depicted in (***a***). Note that the value for the lagging-end overhangs is predicted to plateau (at ∼205 nt) while the leading-end overhangs are constant at 97 nt. Therefore, the average overhang length is predicted to plateau at ∼150 nt which represents a doubling of the length (starting length 74 nt before CTC1 deletion) consistent with the two-fold increase in the relative overhang signal observed in Fig. 3c.

**Extended Data Figure 8.**
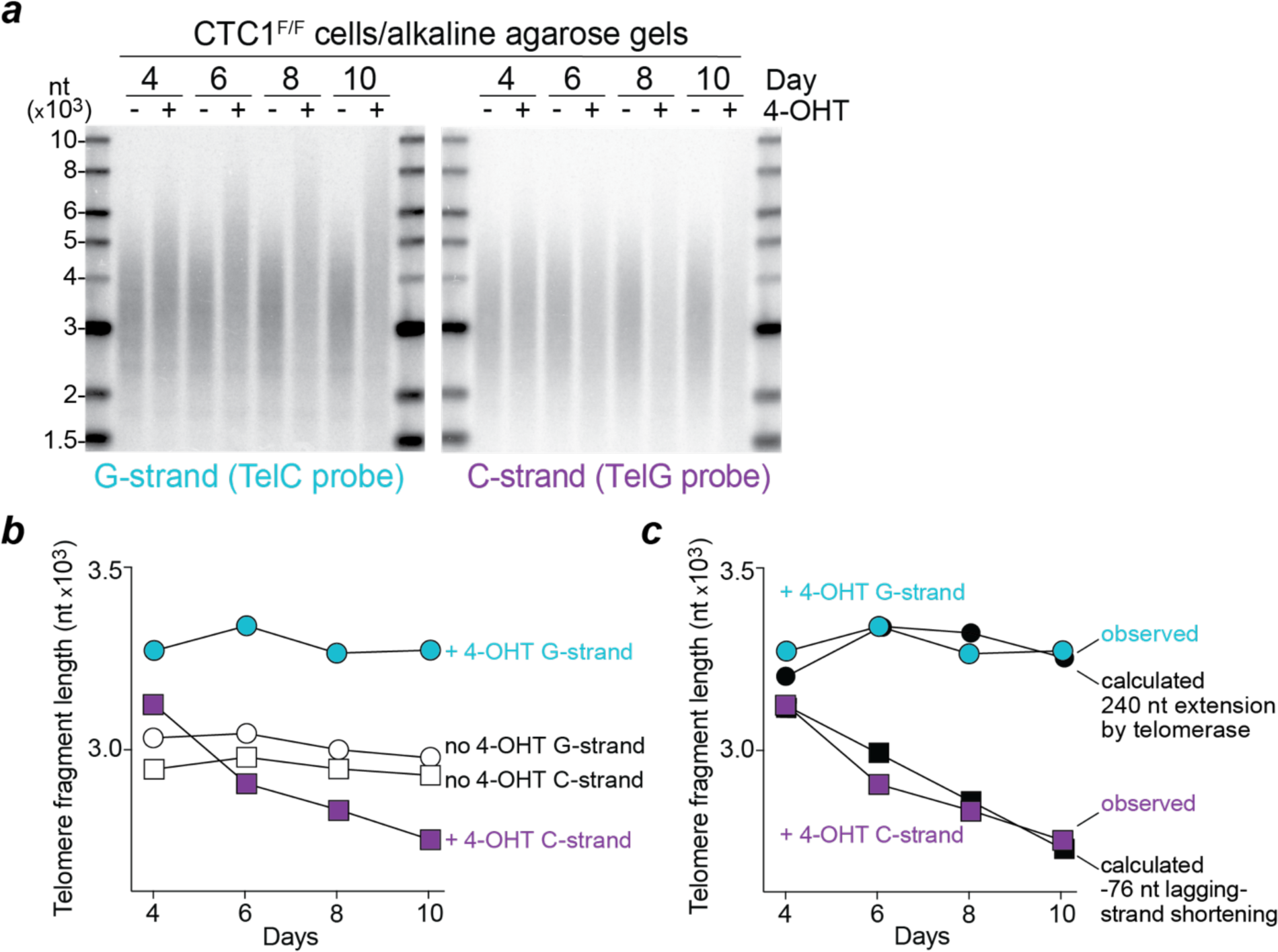
C-strand shortening in HCT116 cells lacking Ctc1. ***a-c***, Measurement of C-strand shortening CTC1^F/F^ cells with or without 4-OHT treatment based on alkaline gel analysis of the telomeric G- and C-strands as in Fig. 4b-d.

**Extended Data Figure 9.**
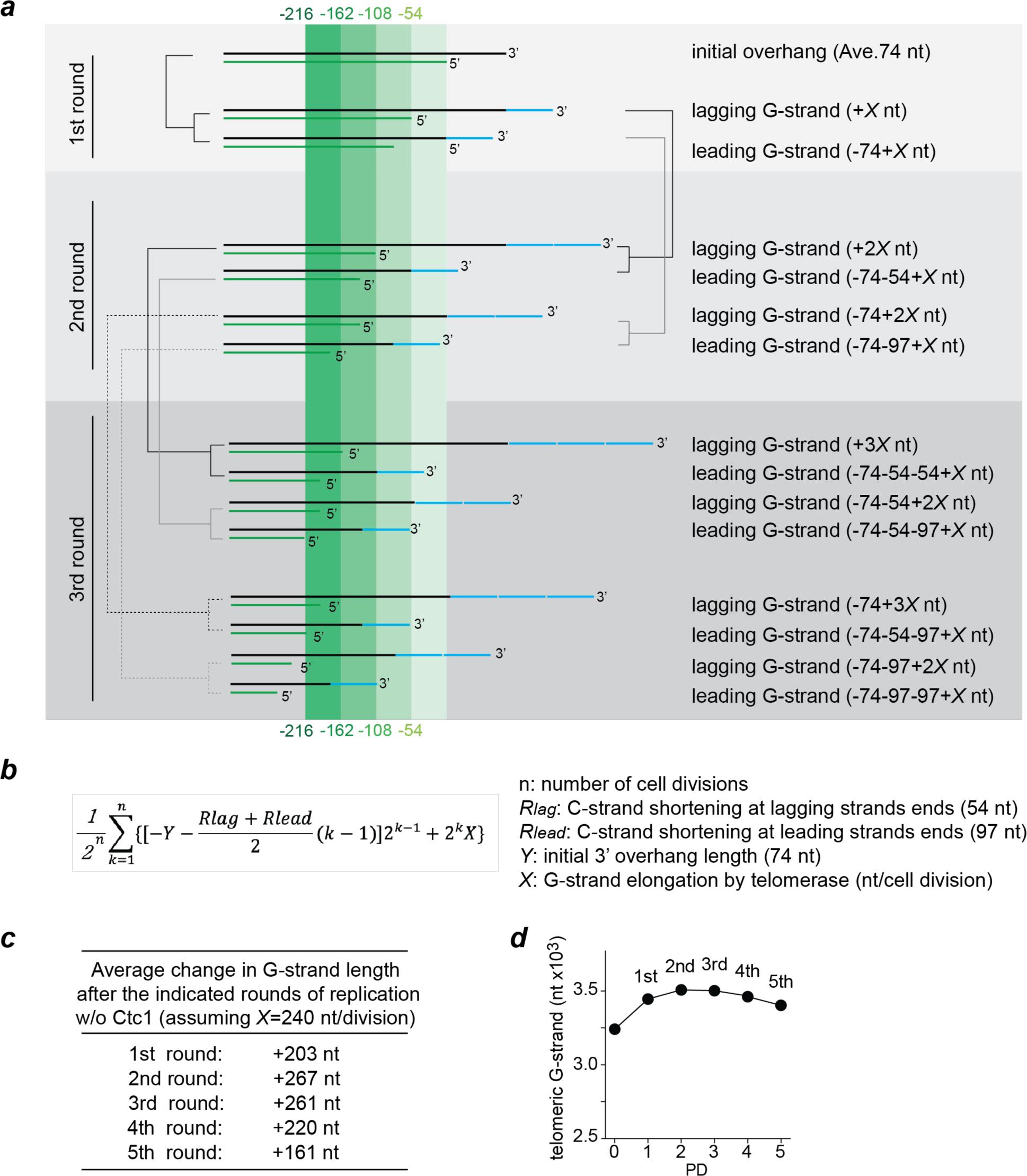
Schematic illustrating the dynamics of the C- and G-strand in Ctc1-deficient cells expressing telomerase. ***a***, Schematic showing the changes of G- and C-strand telomere length over three rounds of replication in telomerase-proficient cells lacking Ctc1. G-strand sequences added by telomerases are depicted with a blue line. The numbers given are as follows (based on Fig. 2 ***j****, **k*** and Extended Data Fig. 6). *X*: nucleotides added by telomerase; 74 nt: initial 3ʹ overhang length; 54 nt: loss of sequences due to incomplete lagging-strand synthesis; 97 nt: resection of the C-strand at leading-end telomeres. ***b***, Equation describing changes in average G-strand length after each round of replication (n) relative to initial G-strand telomere length. ***c***, Increase in average G-strand length with cell divisions calculated using the equation in (***b***) and assuming that telomerase adds an average of 240 nt (*X*) to each end. ***d***, Modeled changes in the length of the G-strand in Ctc1-deficient cells based on the initial G-strand length of 3.24 kb. To determine *X*, the value for telomerase extension was increased stepwise by 10 nt, plotted and the plot was compared to the actual G-strand changes obtained in Fig. 4d. When *X* is 240 nt, the model plot is a close fit with the *in vivo* data as shown in Fig. 4d.

## References

1. Greider, C. W. & Blackburn, E. H. Identification of a specific telomere terminal transferase activity in Tetrahymena extracts. Cell 43, 405–413 (1985).

2. Lingner, J., Cooper, J. P. & Cech, T. R. Telomerase and DNA end replication: no longer a lagging strand problem. Science 269, 1533–1534 (1995).

3. Griffith, J. D., Comeau, L., Rosenfield, S., Stansel, R. M., Bianchi, A., Moss, H. & de Lange, T. Mammalian telomeres end in a large duplex loop. Cell 97, 503–514 (1999).

4. Doksani, Y., Wu, J. Y., de Lange, T. & Zhuang, X. Super-resolution fluorescence imaging of telomeres reveals TRF2-dependent T-loop formation. Cell 155, 345–356 (2013).

5. Watson, J. D. Origin of concatemeric T7 DNA. Nat New Biol 239, 197–201 (1972).

6. Olovnikov, A. M. A theory of marginotomy. The incomplete copying of template margin in enzymic synthesis of polynucleotides and biological significance of the phenomenon. J Theor Biol 41, 181–190 (1973).

7. Greider, C. W. & Blackburn, E. H. Telomeres, telomerase and cancer. Sci Am 274, 92–97 (1996).

8. Zhao, Y., Sfeir, A. J., Zou, Y., Buseman, C. M., Chow, T. T., Shay, J. W. & Wright, W. E. Telomere extension occurs at most chromosome ends and is uncoupled from fill-in in human cancer cells. Cell 138, 463–475 (2009).

9. Hirai, Y., Masutomi, K. & Ishikawa, F. Kinetics of DNA replication and telomerase reaction at a single-seeded telomere in human cells. Genes Cells 17, 186–204 (2012).

10. Wu, P., Takai, H. & de Lange, T. Telomeric 3’ overhangs derive from resection by Exo1 and Apollo and fill-in by POT1b-associated CST. Cell 150, 39–52 (2012).

11. Wang, F., Stewart, J. A., Kasbek, C., Zhao, Y., Wright, W. E. & Price, C. M. Human CST has independent functions during telomere duplex replication and C-strand fill-in. Cell Rep 2, 1096–1103 (2012).

12. Dai, X., Huang, C., Bhusari, A., Sampathi, S., Schubert, K. & Chai, W. Molecular steps of G-overhang generation at human telomeres and its function in chromosome end protection. EMBO J 29, 2788–2801 (2010).

13. Stewart, J. A., Wang, Y., Ackerson, S. M. & Schuck, P. L. Emerging roles of CST in maintaining genome stability and human disease. Front Biosci (Landmark Ed) 23, 1564–1586 (2018).

14. Chow, T. T., Zhao, Y., Mak, S. S., Shay, J. W. & Wright, W. E. Early and late steps in telomere overhang processing in normal human cells: the position of the final RNA primer drives telomere shortening. Genes Dev 26, 1167–1178 (2012).

15. Yeeles, J. T., Deegan, T. D., Janska, A., Early, A. & Diffley, J. F. Regulated eukaryotic DNA replication origin firing with purified proteins. Nature 519, 431–435 (2015).

16. Yeeles, J. T. P., Janska, A., Early, A. & Diffley, J. F. X. How the Eukaryotic Replisome Achieves Rapid and Efficient DNA Replication. Mol Cell 65, 105–116 (2017).

17. Soudet, J., Jolivet, P. & Teixeira, M. T. Elucidation of the DNA end-replication problem in Saccharomyces cerevisiae. Mol Cell 53, 954–964 (2014).

18. Aria, V. & Yeeles, J. T. P. Mechanism of Bidirectional Leading-Strand Synthesis Establishment at Eukaryotic DNA Replication Origins. Mol Cell 73, 199–211 (2019).

19. Jones, M. L., Aria, V., Baris, Y. & Yeeles, J. T. P. How Pol α-primase is targeted to replisomes to prime eukaryotic DNA replication. Mol Cell 83, 2911–2924.e16 (2023).

20. Lim, C. J. & Cech, T. R. Shaping human telomeres: from shelterin and CST complexes to telomeric chromatin organization. Nat Rev Mol Cell Biol 22, 283–298 (2021).

21. Cai, S. W. & de Lange, T. CST-Polα/Primase: the second telomere maintenance machine. Genes Dev 37, 555–569 (2023).

22. Goulian, M., Heard, C. J. & Grimm, S. L. Purification and properties of an accessory protein for DNA polymerase alpha/primase. J Biol Chem 265, 13221–13230 (1990).

23. Goulian, M. & Heard, C. J. The mechanism of action of an accessory protein for DNA polymerase alpha/primase. J Biol Chem 265, 13231–13239 (1990).

24. Lim, C. J., Barbour, A. T., Zaug, A. J., Goodrich, K. J., McKay, A. E., Wuttke, D. S. & Cech, T. R. The structure of human CST reveals a decameric assembly bound to telomeric DNA. Science 368, 1081–1085 (2020).

25. Lue, N. F. Evolving Linear Chromosomes and Telomeres: A C-Strand-Centric View. Trends Biochem Sci 43, 314–326 (2018).

26. Gao, H., Cervantes, R. B., Mandell, E. K., Otero, J. H. & Lundblad, V. RPA-like proteins mediate yeast telomere function. Nat Struct Mol Biol 14, 208–214 (2007).

27. Miyake, Y., Nakamura, M., Nabetani, A., Shimamura, S., Tamura, M., Yonehara, S., Saito, M. & Ishikawa, F. RPA-like mammalian Ctc1-Stn1-Ten1 complex binds to single-stranded DNA and protects telomeres independently of the Pot1 pathway. Mol Cell 36, 193–206 (2009).

28. Surovtseva, Y. V., Churikov, D., Boltz, K. A., Song, X., Lamb, J. C., Warrington, R., Leehy, K., Heacock, M., Price, C. M. & Shippen, D. E. Conserved telomere maintenance component 1 interacts with STN1 and maintains chromosome ends in higher eukaryotes. Mol Cell 36, 207–218 (2009).

29. Cai, S. W., Zinder, J. C., Svetlov, V., Bush, M. W., Nudler, E., Walz, T. & de Lange, T. Cryo-EM structure of the human CST-Polα/primase complex in a recruitment state. Nat Struct Mol Biol 29, 813–819 (2022).

30. He, Q., Lin, X., Chavez, B. L., Agrawal, S., Lusk, B. L. & Lim, C. J. Structures of the human CST-Polα-primase complex bound to telomere templates. Nature 608, 826–832 (2022).

31. Zaug, A. J., Goodrich, K. J., Song, J. J., Sullivan, A. E. & Cech, T. R. Reconstitution of a telomeric replicon organized by CST. Nature 608, 819–825 (2022).

32. Feng, X., Hsu, S. J., Kasbek, C., Chaiken, M. & Price, C. M. CTC1-mediated C-strand fill-in is an essential step in telomere length maintenance. Nucleic Acids Res 45, 4281–4293 (2017).

33. Takai, H., Jenkinson, E., Kabir, S., Babul-Hirji, R., Najm-Tehrani, N., Chitayat, D. A., Crow, Y. J. & de Lange, T. A POT1 mutation implicates defective telomere end fill-in and telomere truncations in Coats plus. Genes Dev 30, 812–826 (2016).

34. McElligott, R. & Wellinger, R. J. The terminal DNA structure of mammalian chromosomes. EMBO J 16, 3705–3714 (1997).

35. Damm, K., Hemmann, U., Garin-Chesa, P., Hauel, N., Kauffmann, I., Priepke, H., Niestroj, C., Daiber, C., Enenkel, B., Guilliard, B., Lauritsch, I., Müller, E., Pascolo, E., Sauter, G., Pantic, M., Martens, U. M., Wenz, C., Lingner, J., Kraut, N., Rettig, W. J. & Schnapp, A. A highly selective telomerase inhibitor limiting human cancer cell proliferation. EMBO J 20, 6958–6968 (2001).

36. Larrivée, M., LeBel, C. & Wellinger, R. J. The generation of proper constitutive G-tails on yeast telomeres is dependent on the MRX complex. Genes Dev 18, 1391–1396 (2004).

37. Baird, D. M., Rowson, J., Wynford-Thomas, D. & Kipling, D. Extensive allelic variation and ultrashort telomeres in senescent human cells. Nat Genet 33, 203–207 (2003).

38. Tesmer, V. M., Brenner, K. A. & Nandakumar, J. Human POT1 protects the telomeric ds-ss DNA junction by capping the 5’ end of the chromosome. Science 381, 771–778 (2023).

39. Hockemeyer, D., Daniels, J. P., Takai, H. & de Lange, T. Recent expansion of the telomeric complex in rodents: Two distinct POT1 proteins protect mouse telomeres. Cell 126, 63–77 (2006).

40. Anderson, B. H., Kasher, P. R., Mayer, J., Szynkiewicz, M., Jenkinson, E. M., Bhaskar, S. S., Urquhart, J. E., Daly, S. B., Dickerson, J. E., O’Sullivan, J., Leibundgut, E. O., Muter, J., Abdel-Salem, G. M., Babul-Hirji, R., Baxter, P., Berger, A., Bonafé, L., Brunstom-Hernandez, J. E., Buckard, J. A., Chitayat, D., Chong, W. K., Cordelli, D. M., Ferreira, P., Fluss, J., Forrest, E. H., Franzoni, E., Garone, C., Hammans, S. R., Houge, G., Hughes, I., Jacquemont, S., Jeannet, P. Y., Jefferson, R. J., Kumar, R., Kutschke, G., Lundberg, S., Lourenço, C. M., Mehta, R., Naidu, S., Nischal, K. K., Nunes, L., Ounap, K., Philippart, M., Prabhakar, P., Risen, S. R., Schiffmann, R., Soh, C., Stephenson, J. B., Stewart, H., Stone, J., Tolmie, J. L., van der Knaap, M. S., Vieira, J. P., Vilain, C. N., Wakeling, E. L., Wermenbol, V., Whitney, A., Lovell, S. C., Meyer, S., Livingston, J. H., Baerlocher, G. M., Black, G. C., Rice, G. I. & Crow, Y. J. Mutations in CTC1, encoding conserved telomere maintenance component 1, cause Coats plus. Nat Genet 44, 338–342 (2012).

41. Komura, J. & Riggs, A. D. Terminal transferase-dependent PCR: a versatile and sensitive method for in vivo footprinting and detection of DNA adducts. Nucleic Acids Res 26, 1807–1811 (1998).

42. Hanish, J. P., Yanowitz, J. L. & de Lange, T. Stringent sequence requirements for the formation of human telomeres. Proc Natl Acad Sci U S A 91, 8861–8865 (1994).

43. Smogorzewska, A., Karlseder, J., Holtgreve-Grez, H., Jauch, A. & de Lange, T. DNA ligase IV-dependent NHEJ of deprotected mammalian telomeres in G1 and G2. Curr Biol 12, 1635–1644 (2002).

44. Harley, C. B., Futcher, A. B. & Greider, C. W. Telomeres shorten during ageing of human fibroblasts. Nature 345, 458–460 (1990).

